# Axenic *in-vitro* cultivation of nineteen peat-moss (*Sphagnum* L.) species as a resource for basic biology, biotechnology and paludiculture

**DOI:** 10.1101/2020.03.25.004770

**Authors:** Melanie A. Heck, Volker M. Lüth, Matthias Krebs, Mira Kohl, Anja Prager, Hans Joosten, Eva L. Decker, Ralf Reski

## Abstract

- The cultivation of *Sphagnum* mosses reduces CO_2_ emissions by rewetting drained peatlands and by substituting peat with renewable biomass. ‘Sphagnum farming’ requires large volumes of founder material, which can only be supplied sustainably by axenic cultivation in bioreactors.
- We established axenic *in-vitro* cultures from sporophytes of 19 *Sphagnum* species collected in Austria, Germany, Latvia, Netherlands, Russia and Sweden, namely *S. angustifolium, S. balticum, S. capillifolium, S. centrale, S. compactum, S. cuspidatum, S. fallax, S. fimbriatum, S. fuscum, S. lindbergii, S. medium/divinum, S. palustre, S. papillosum, S. rubellum, S. russowii, S. squarrosum, S. subnitens, S. subfulvum*, and *S. warnstorfii*. These species cover five of the six European *Sphagnum* sections, namely *Acutifolia, Cuspidata, Rigida, Sphagnum* and *Squarrosa*.
- Their growth was measured in axenic suspension cultures, whereas their ploidy was determined by flow cytometry and compared with the genome size of *Physcomitrella patens*. We identified haploid and diploid *Sphagnum* species, found that their cells are predominantly arrested in the G1-phase of the cell cycle, and did not find a correlation between plant productivity and ploidy.
- With this collection, high-quality founder material for diverse large-scale applications but also for basic *Sphagnum* research is available from the International Moss Stock Center (IMSC).

## Introduction

Peatlands cover over four million square kilometres, comprising 3% of Earth’s land and freshwater surface (Joosten & Clarke, 2002), and contain about 30% of the global soil carbon (Gorham, 1991; Frolking and Roulet, 2007). Most peatlands in temperate and boreal zones were formed and are dominated by peat mosses, i.e. mosses of the genus *Sphagnum* (Clymo & Hayward, 1982; Joosten *et al*., 2017). They accumulate dead organic matter (‘peat’) under wet, anoxic, acidic and nutrient-poor conditions, which lower microbial activity and reduce the decay of organic matter. As a result, pristine peatlands are carbon sinks, i.e. they sequester more carbon than they emit, and function as long-term carbon stores (Clymo & Hayward, 1982; Joosten *et al*., 2016). Climate change scenarios assume that prolonged droughts, elevated temperatures and increased nitrogen deposition (Galloway *et al*., 2008) decrease the growth of *Sphagnum* mosses and increase decay, thus reducing the amount of sequestered carbon (Limpens *et al*., 2011; Norby *et al*., 2019). Moreover, changing microbial communities might enhance the functional shift from sink to source (Lew *et al*., 2019; Juan-Ovejero *et al*., 2020; Rewcastle *et al*., 2020). Together, its impact on global carbon cycling makes *Sphagnum* an important ecological model, attracting a growing number of scientists. Consequently, the first draft genome of *Sphagnum fallax* became available recently (v0.5, http://phytozome.jgi.doe.gov/) (Weston *et al*., 2018). Although *Sphagnum* mosses are of growing economic importance for many applications, including waste water treatment (Couillard, 1994), as sensors of air pollution (Capozzi *et al*., 2016, 2017; Di Palma *et al*., 2019; Aboal *et al*., 2020) and as raw material for growing media (Wichmann et al. 2020), they are not yet analysed in great detail.

The global area of peatlands has been reduced significantly (10 to 20%) since 1800, in particular by drainage for agriculture and forestry. Moreover, peat serves as energy generation, and as substrate for horticulture (Joosten & Clarke, 2002). Drainage leads to peat mineralization and subsequent emissions of greenhouse gases (GHGs), such as CO_2_ and N_2_O (Van Den Pol-Van Dasselaar *et al*., 1999; Boon *et al*., 2014; Carlson *et al*., 2017). While drained peatlands cover only 0.4% of the land surface, they are responsible for 32% of cropland and almost 5% of anthropogenic GHG emissions globally (Joosten *et al*., 2016; Carlson *et al*., 2017). Leifeld *et al*. (2019) estimated that in 1960 the global peatland biome turned from a net sink to a net source of soil-derived GHGs. Further, these authors predict a cumulative emission from drained peatlands of 249 +/- 38 petagrams of CO_2_ equivalent by 2100 if the current trend continues.

Rewetting of drained peatlands decreases these emissions and may even restore the carbon sink function (Joosten *et al*., 2016; Wichtmann *et al*., 2016). Rewetting, however, makes conventional drainage-based land use impossible (Wichmann *et al*., 2017). Paludiculture, wet agriculture and forestry on peatlands, allows land use to continue and to combine emission reduction with biomass production. It includes traditional peatland cultivation (reed mowing, litter usage) and new approaches for utilization (Abel *et al*., 2013; Wichtmann *et al*., 2016).

Sphagnum farming on rewetted bogs is a promising example of paludiculture as it produces *Sphagnum* biomass as a substitute for peat (Gaudig & Joosten, 2002; Gaudig *et al*., 2014, 2018; Decker & Reski, 2020). It decreases GHG emissions substantially by rewetting drained peatlands, by avoiding the use and oxidation of fossil peat, and by preserving hitherto undrained peatlands as carbon stores and sinks (Wichtmann *et al*., 2016; Günther *et al*., 2017). Potential sites for Sphagnum farming are degraded bogs and acidic water bodies (Wichmann *et al*., 2017).

Different environmental conditions, e.g. water level or nutrient supply, and different requirements of the produced biomass call for a variety of peat-moss species and genotypes as founder material for Sphagnum farms (Gaudig *et al*., 2018). Lack of sufficient founder material is currently a major bottleneck for the large-scale implementation of Sphagnum farming. In the European Union, *Sphagnum* species and their habitats are protected by the Council Directive 92/43/EEC, constraining the collection of founder material from natural habitats. Furthermore, commercial Sphagnum farming requires *Sphagnum* material without unwanted biological contaminations (Gaudig *et al*., 2018) and of a constitution that is fit-for-purpose. Moss clones established from a single spore or plant share the same genetic, physiological and environmental background, allowing the multiplication of selected clones to achieve maximum yields. *Sphagnum* founder material of controlled quality can be produced under aseptic conditions with standard tissue culture methods (Caporn *et al*., 2017), but probably more rapidly by axenic cultivation in bioreactors.

An important step towards the large-scale production of such founder material was the development of an axenic photobioreactor production process for *Sphagnum palustre*, using monoclonal material generated from a single spore (Beike *et al*., 2015). Under standardized laboratory conditions, the multiplication rate of the material was up to 30-fold within four weeks. The phenotypic characteristics of *in-vitro* cultivated *S. palustre* plants were comparable to those of plants from natural habitats (Beike *et al*., 2015). A further improvement to vegetative *in-vitro* propagation of *Sphagnum* is the establishment of a protonema-proliferation protocol for *S. squarrosum* (Zhao *et al*., 2019). Besides these two species, we found only one report of the establishment of *S. fallax* cultures in a bioreactor (Rudolph *et al*., 1988) and to our knowledge, no other *Sphagnum* species were analysed in axenic laboratory conditions so far.

Here, we report on the establishment of axenic *in-vitro* cultures of 19 *Sphagnum* species from five peat-moss sections and compare their growth behaviour, their genome size and their cell cycle. With this collection, high-quality founder material for diverse large-scale applications, but also for basic *Sphagnum* research is available.

## Materials & methods

### Decontamination of sporophytes and spore germination

We collected sporophytes from 19 *Sphagnum* species in the field (Table 1) and stored them at 4°C. The taxonomic status of the *S. medium/divinum* clones needs clarification to accommodate for taxonomic insights (Hassel *et al*., 2018) after collection. Spore capsules were surface-sterilized and opened with forceps in 1 ml of sodium hypochlorite solution. The solution was freshly prepared with autoclaved water with 2 drops of Tween 20 per 500 ml H_2_O and a final concentration of either 0.6%, 1.2% or 2.4% NaClO. The incubation was stopped at different time points between 30 sec and 7 min by transferring 100 μl of the suspension to 1 ml autoclaved water. From this dilution, 500 μl were transferred to a sterile Petri dish, which contained one of the following solid media: 1) Knop medium (1.84 mM KH2PO4, 3.35 mM KCl, 1.01 mM MgSO_4_, 4.24 mM Ca(NO_3_)_2_, 45 μM FeSO_4_) according to Reski & Abel (1985) supplemented with microelements (50 μM H_3_BO_3_, 50 μM MnSO_4_, 15 μM ZnSO_4_, 2.5 μM KJ, 500 nM Na_2_MoO_4_, 50 nM CuSO_4_, 50 nM Co(NO_3_)_2_) according to Schween *et al*. (2003a), or 2) *Sphagnum* medium (Knop medium with microelements (ME), 0.3 % sucrose and 1.25 mM NH_4_NO_3_) according to Beike *et al*. (2015). Petri dishes were sealed with Parafilm (Carl Roth, Germany), and cultivated under standard growth conditions: climate chamber, temperature of 22°C, photoperiod regime of 16 h : 8 h (light : dark), light intensity of 70±5 μmol m^−2^ s^−1^ provided by fluorescent tubes. Light intensities were measured with a planar quantum sensor (Li-Cor 250, Li-Cor Biosciences, Bad Homburg, Germany).

After spore germination, filaments were separated and transferred to new Petri dishes containing either solid Knop ME or solid Sphagnum medium under sterile conditions using needles and a stereo microscope (Stemi 2000-C, Zeiss, Jena, Germany). Plates containing one of the following media: 1) Knop ME supplemented with 1% glucose and 12 g l^−1^ Purified Agar (Oxoid Limited, UK), 2) LB (10 g l^−1^ Bacto Tryptone (Becton, Dickinson and Company, NJ, USA), 10 g l^−1^ NaCl, 5 g l^−1^ Bacto Yeast Extract (Becton, Dickinson and Company) and 15 g l^−1^ Bacto Agar (Becton, Dickinson and Company)), or 3) Tryptic Soy Agar (TSA) with 1% glucose (15 g l^−1^ peptone from casein, 5 g l^−1^ soy peptone, 5 g l^−1^ NaCl) and 12 g l^−1^ purified agar (Oxoid Limited, UK) served as controls. These plates were sealed with Parafilm, stored and inspected at room temperature. If no contamination occurred within four weeks, we considered a culture as axenic.

### *In-vitro* cultivation

Gametophores were cultivated on solid media and in suspension. For cultivation on solid medium, gametophores were transferred to Knop ME or Sphagnum medium. The Petri dishes were sealed with Parafilm and cultivated under standard conditions.

For suspension cultures, gametophores were disrupted with forceps in laminar flow benches, and transferred to 35 ml Sphagnum medium in 100 ml Erlenmeyer flasks. Flasks were closed with Silicosen^®^ silicone sponge plugs (Hirschmann Laborgeräte, Eberstadt, Germany) and agitated on a rotary shaker at 120 rpm (B. Braun Biotech International, Melsungen, Germany) under standard conditions.

### Light microscopy

Gametophores were analysed with a stereo microscope (SZX7, Olympus Corporation, Tokyo, Japan) and a camera (AxioCam ICc 1, Zeiss, Jena, Germany). Photographs were scaled with AxioVision software 4.8 (Zeiss). Stacks of images with different focal points were combined with CombineZ 5.3 (Alan Hadley, https://combinezp.software.informer.com/).

### Growth determination

Plant growth was determined on solid media as well as in suspension (**Fig. 1**). First, growth of up to 16 clones of each species was determined on agar plates with both Knop ME and Sphagnum medium. Clones were randomly selected from all available spore capsules. The uppermost five millimetres from the tip of each gametophore (the capitulum) was cut, transferred to solid media and cultivated under standard conditions for four weeks (**Fig. 1**). Growth was documented photographically every week. The pictures were transferred into binary images and the area was assessed by counting the pixels (**Fig. 1a**) using ImageJ version 1.51f (Wayne Rasband, https://imagej.nih.gov/ij/). Only the area of growth was analysed, not the height of the gametophore. Additionally, shape and colour were visually assessed to select the six largest clones of each species after four weeks of growth. These were subsequently assessed for biomass increase in suspensions.

**Fig. 1.**
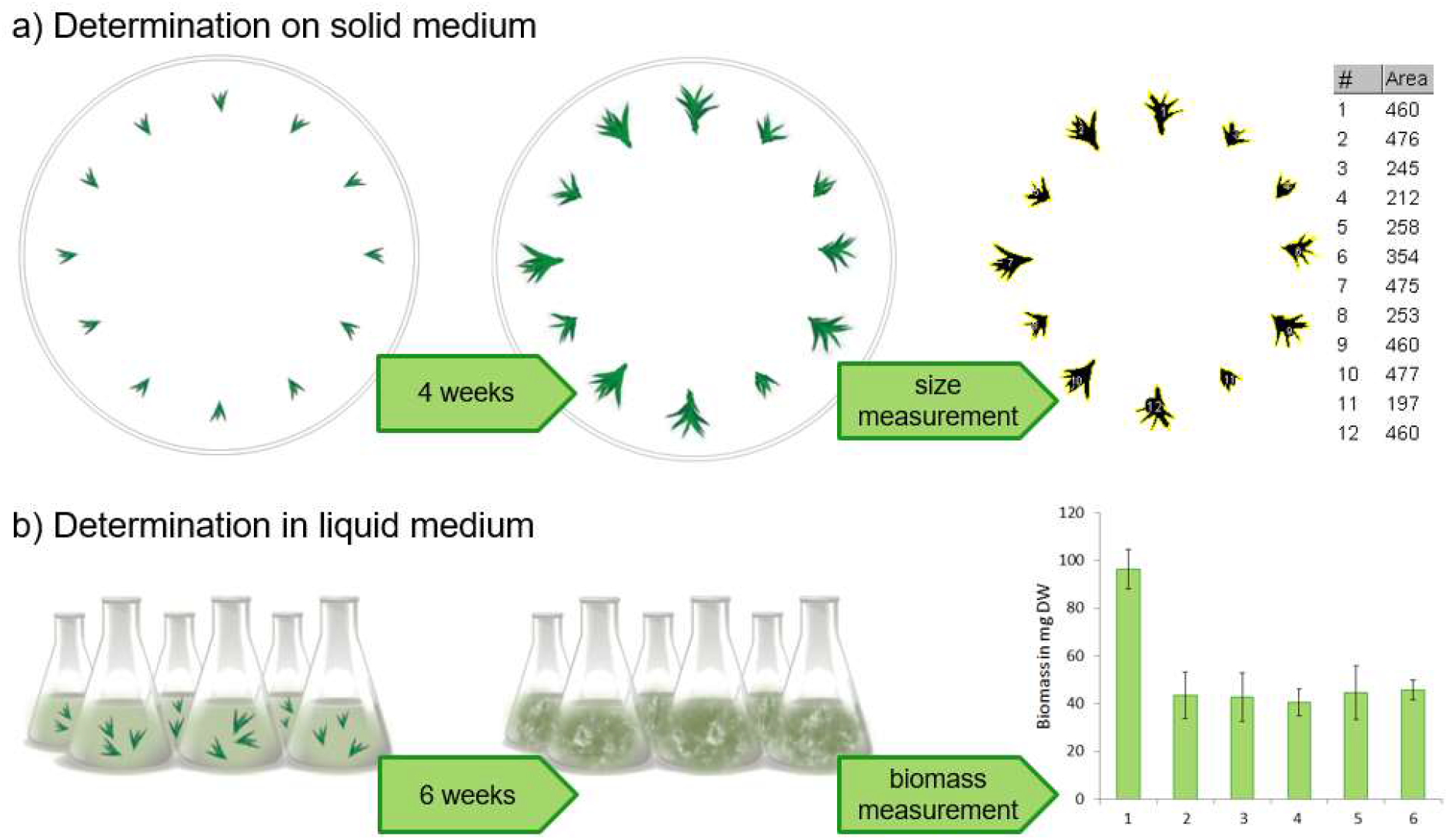
Schematic representation of the growth determination of *Sphagnum* spp. a) on solid medium, b) in suspension cultures. a) Gametophores of the same size were transferred to a Petri dish and cultivated for four weeks. The growth was documented photographically and the size analysed by image processing supported area measurement using ImageJ. b) Gametophores were transferred to Erlenmeyer flasks and cultivated for six weeks. Growth was determined by dry weight measurement of the biomass.

Three gametophores per clone were transferred to 50 ml Sphagnum medium in 100 ml Erlenmeyer flasks and cultivated under standard conditions for six weeks. Subsequently, the total biomass was harvested by filtering with a Büchner funnel and a vacuum pump. The moss material was transferred to pre-dried (0.5 h at 105°C) aluminium weighing pans (Köhler Technische Produkte, Neulußheim, Germany) and dried for 2 hours at 105°C. Subsequently, the dry weight (DW) was determined with an accuracy scale (CPA 3245, Sartorius, Göttingen, Germany) (**Fig. 1b**). The clone with the highest DW increase of each species was selected as best grown clone.

### Flow-cytometry

Ploidy levels of the six best-grown clones of each species were determined via flow-cytometry (FCM). Gametophores were chopped with a razor blade in 0.5 ml 4’,6-diamidino-2-phenylindol (DAPI) solution (Carl Roth, Germany) containing 0.01 mg l^−1^ DAPI, 1.07 g l-^1^ MgCl_2_·6H_2_O, 5 g l^−1^ NaCl, 21.11 g l^−1^ TRIS and 1 ml l^−1^ Triton X-100. Afterwards, 1.5 ml DAPI solution was added and the material filtered through a 30μm sieve, and subsequently analysed with a Flow Cytometer Partec CyFlow^®^ Space (Sysmex Partec, Görlitz, Germany), equipped with a 365 nm UV-LED. *Physcomitrella patens* protonema served as internal standard (modified after Schween *et al*., 2003b).

### Statistical analysis

To determine significance values between the growths of the clones, data were analysed by one-way analysis of variance (ANOVA), where p values below 0.05 were considered as significant. Afterwards, each data set was tested with Student’s t-test, where ***, ** and * denote significance at the 0.1%, 1% and 5% level, respectively. Statistical analyses were performed with GraphPad Prism^®^ and diagrams were created with Excel 2016.

## Results and discussion

### Induction of axenic *in-vitro* cultures

Surface sterilization of spore capsules is an established method to start axenic *in-vitro* cultivation (Beike *et al*., 2015). *Sphagnum* spores are still viable after 13 years when stored in the cold and they can form persistent spore banks in nature (Sundberg & Rydin, 2000). We observed spore germination in *S. angustifolium* and *S. fimbriatum* after three years storage at 4°C. However, we decontaminated most sporophytes within two months after collection. Detergent concentration and exposure time were adjusted individually for each sporophyte. We did not find a correlation between species or sporophyte maturation level and exposure time for successful decontamination and germination of spores. Sugar accelerated spore germination of all species except *S. compactum*, but all species except *S. warnstorfii* also germinated on Knop ME. Filaments developed from sterilized spores after 2 – 20 weeks with high variations within every species. Single gametophores were separated, and subsequently cultivated on solid medium as independent clones.

Beike *et al*. (2015) showed that *in-vitro* cultivated *S. palustre* plants and plants taken from natural habitats have similar phenotypic characteristics. However, in that study *in-vitro* grown gametophores were smaller and the shoots had more lanceolate leaves compared to the cucullate, ovate leaves of the thicker and heavier field shoots. We observed such deviating morphological characteristics for all 19 *in-vitro* cultivated *Sphagnum* species (**Fig. 2**). Differences in spore germination, plant development and plant morphology between axenic moss cultures and field-grown mosses may be due, besides obvious abiotic factors and speed of growth, to effects of the microbiome present only in the latter. Cross-kingdom and cross-clade signalling via small molecules can influence morphology of *Sphagnum* similar to *Physcomitrella patens* (Kostka *et al*., 2016; Decker *et al*., 2017; Vesty *et al*., 2020).

**Fig. 2.**
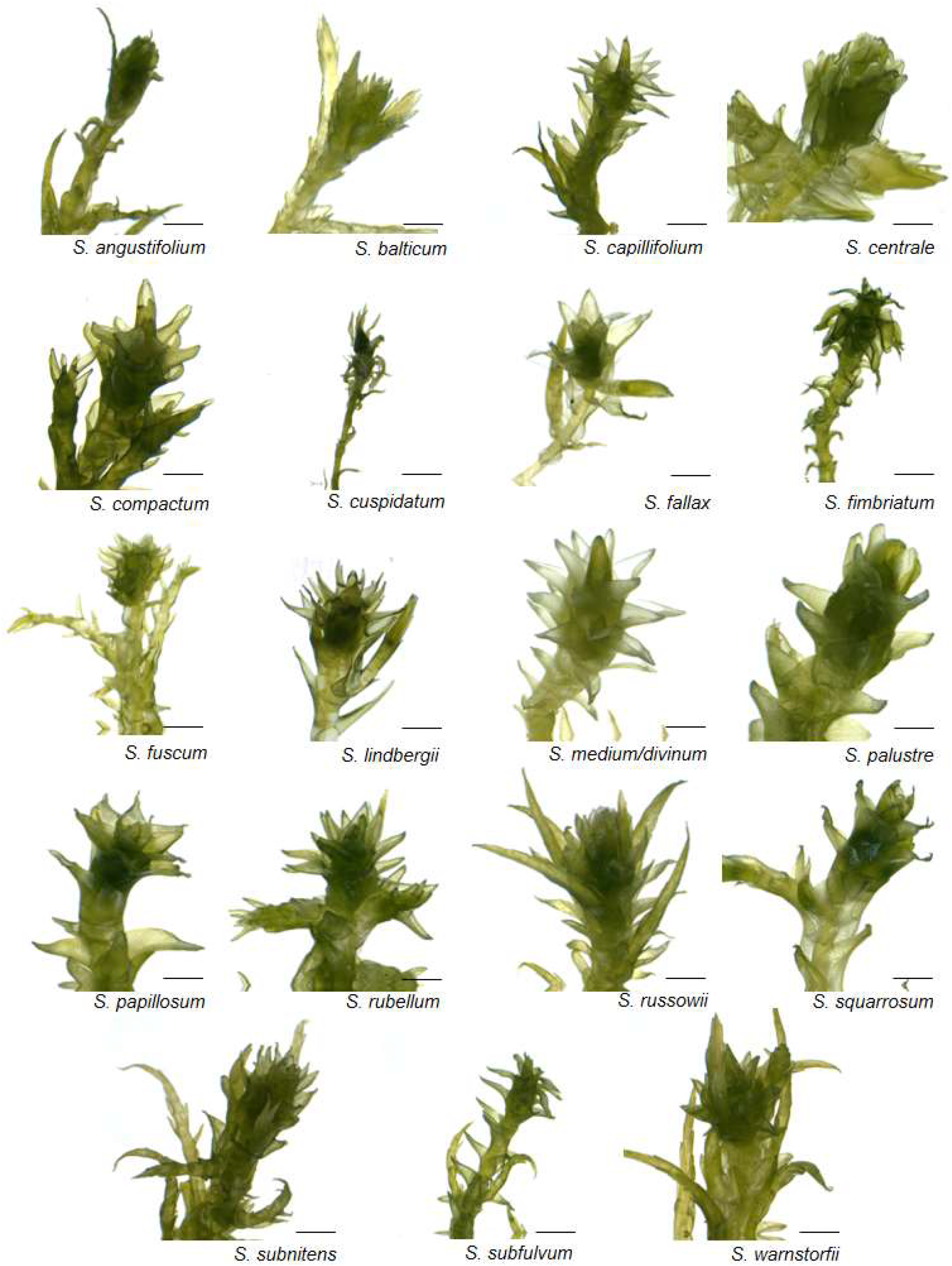
Light microscopic images of characteristic gametophores of *Sphagnum* spp. after four weeks of axenic cultivation on solid Sphagnum medium. Scale bar = 1 mm.

### Selection of the best-growing clones

For subsequent analyses, we reduced the number of clones by preselection on solid medium. Cultivation on solid medium allows long-term storage; while suspension cultures yield higher amounts of biomass (Beike *et al*., 2015).

We describe the selection of the best-growing clone here in detail for *Sphagnum fuscum*, whilst descriptions for the other species are in the supplement (Figures S1–S17). Capitula of eight *S. fuscum* clones were cultivated on solid Knop ME and on solid Sphagnum medium, respectively (**Fig. 3**). Two clones were selected from capsule 1, four clones from capsule 2 and two clones from capsule 3, all collected from the same location in Sweden.

**Fig. 3.**
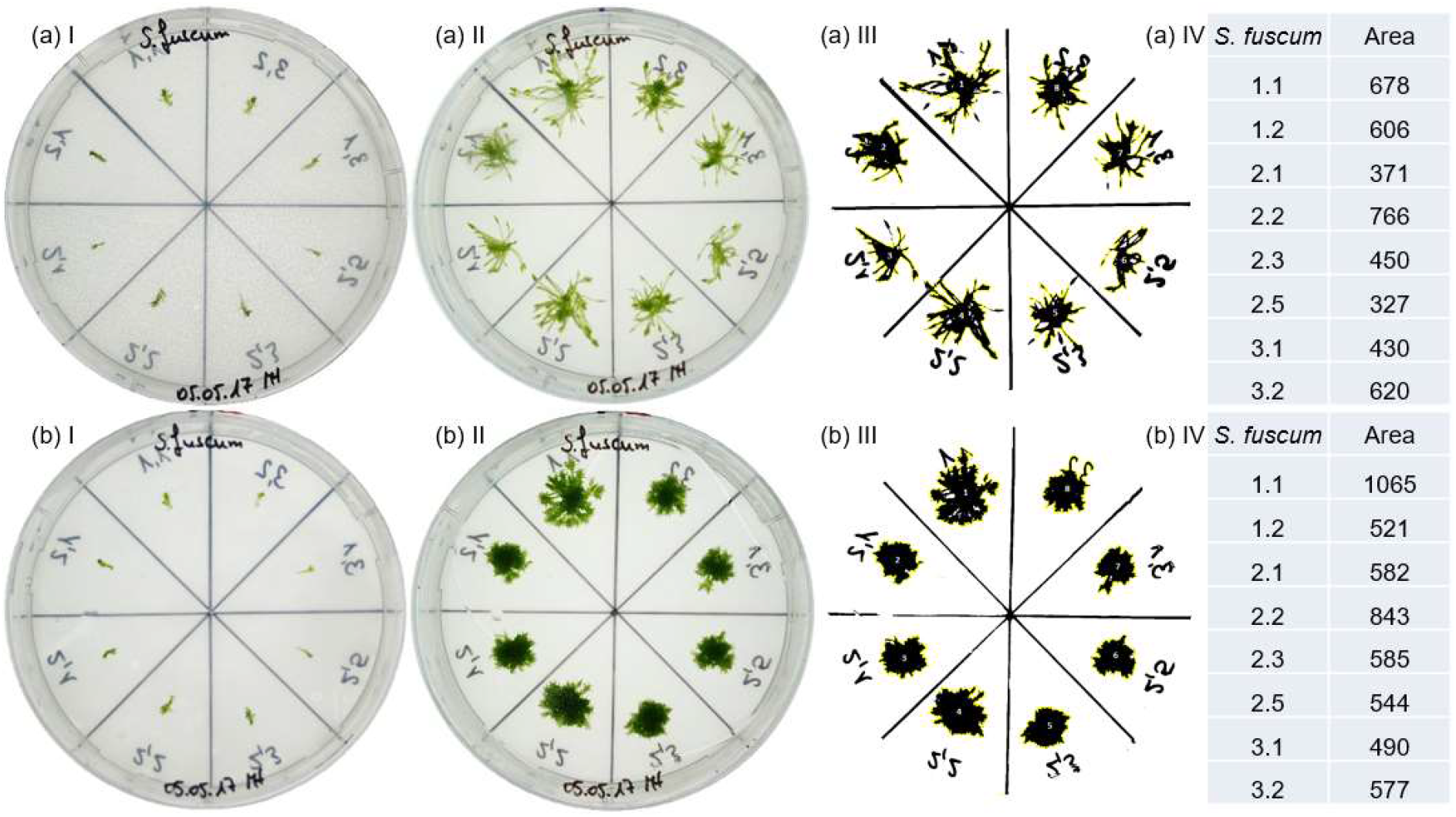
Growth determination of *Sphagnum fuscum* on a) solid Knop ME and b) solid Sphagnum medium. I) Capitula of eight independent clones were cut to 5 mm size and transferred to Petri dishes. II) Gametophores after four weeks of cultivation. III) The size of the gametophores was measured by counting the pixels on binary pictures using ImageJ. IV) The area (number of pixels) of each gametophore.

Sphagnum medium comprises Knop ME, sucrose and ammonium nitrate. Previous studies described growth enhancement of *Sphagnum* by sucrose or other saccharides (Simola, 1969; Graham *et al*., 2010; Beike *et al*., 2015), or a nitrogen source (Simola, 1975; Beike *et al*., 2015). Fertilization, especially the addition of nitrogen and phosphorus, can affect the morphology of *Sphagnum* (Fritz *et al*., 2012).

Gametophores on Sphagnum medium were more compact with a darker green colour compared to gametophores on Knop ME, as depicted for *S. fuscum* in **Fig. 3a II, b II**. We found this effect for all *Sphagnum* species in our study.

The six clones covering the largest area were in descending order 2.2, 1.1, 3.2, 1.2, 2.3 and 3.1 on Knop ME (**Fig. 3a IV**) and 1.1, 2.2, 2.3, 2.1, 3.2 and 2.5 on Sphagnum medium (**Fig. 3b IV**). Clones 1.1, 2.2, 2.3 and 3.2 were among the six best clones on both media, clones 1.2, 2.1, 2.5 and 3.1 among the six best clones on one of the plates. In case of ambiguous results, care was taken that at least one clone of each geographical location remained among the six best clones to maintain the highest possible ecotype variation. In this way, clones 1.1, 2.1, 2.2, 2.3, 3.1 and 3.2 were identified as the six best clones on solid media and subsequently analysed in suspension. Here, clone 1.1 yielded significantly more biomass than the other five clones (**Fig. 4**).

**Fig. 4.**
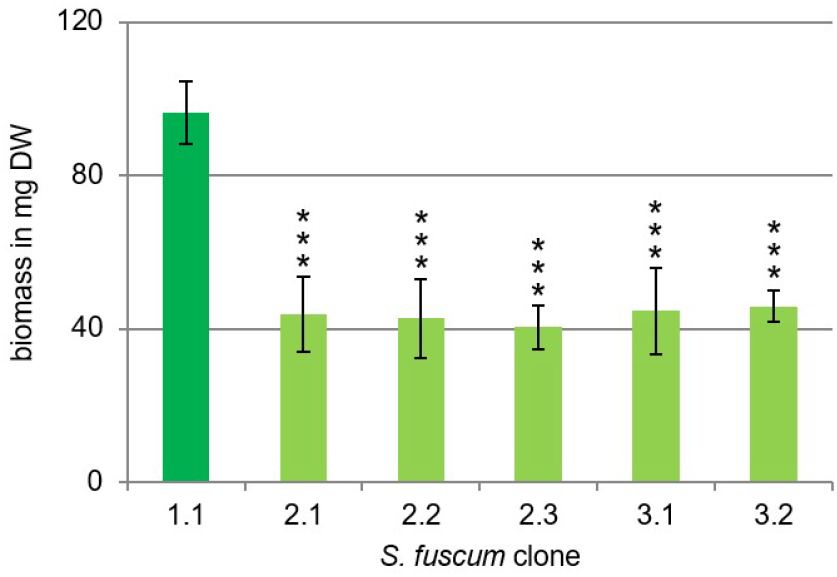
Biomass (in mg dry weight) of six *S. fuscum* clones. The growth of the clones was determined in suspension cultures by measuring the dry weight after cultivation of three capitula in flasks containing 50 ml Sphagnum media for six weeks. The y-axis shows the biomass in dry weight, the x-axis shows the clone. Data represent mean values with standard deviations of three biological replicates (ANOVA p<0.0001). Clone 1.1 yielded significantly more biomass compared to the clones 2.1***, 2.2***, 2.3***, 3.1*** and 3.2***. Asterisks represent results of student t-test performed in comparison to clone 1.1 (*** = p < 0.001).

Up to six best-growing clones per species from different sporophytes were deposited in the International Moss Stock Center (http://www.moss-stock-center.org). All deposited clones, their accession numbers, and the origin of the sporophyte (date of collection, location of collection) are listed in **Table 1**.

The taxonomic status of clones derived from sporophytes originally collected as *S. magellanicum* needs clarification in future because a new species concept of *S. magellanicum, S. medium* and *S. divinum* became available (Hassel *et al*., 2018) during the course of our study. We cannot group these clones into one of these species based on morphology, because it differs slightly between *in-vitro* and the field, as described for *S. palustre* (Beike *et al*., 2015). However, we can distinguish them by their collection site, as geographical distribution differs. Accordingly, our clones most probably are not *S. magellanicum*, because Hassel *et al*. (2018) suggest its occurrence in Argentina and Chile only. In contrast, we collected those sporophytes in Sweden and Russia. Therefore, our clones are most likely *S. divinum* or *S. medium*, because both are circumpolar in the northern hemisphere (Hassel *et al*., 2018). As both species occur in mixed stands (Hassel *et al*., 2018), we are currently not able to separate them by collection site either. Therefore, we list these clones here as *S. medium/divinum*. To resolve this uncertainty in future, a detailed analysis with different molecular markers (Hassel *et al*., 2018; Di Palma *et al*., 2016; von Stackelberg *et al*., 2006) is needed. However, we note that these clones are more heterogeneous regarding morphology, colour and growth rate than clones from the other species in our study. This suggests to the existence of two species in our *S. medium/divinum* collection. If this hypothesis is confirmed by molecular genetic analyses in future, our axenic *Sphagnum* collection comprises 20 instead of 19 species.

**Table 1.** *Sphagnum* spp. clones in axenic culture with corresponding IMSC numbers, date and location of spore capsule collection.

| Species | IMSC No. | 6 best clones | Origin of the spore capsule |  |
| --- | --- | --- | --- | --- |
|  |  |  | Date | Location |
| *S. angustifolium* | 41114 | 2.1, 2.2, 2.3, **2.4**, 2.5, 7.1 | 2015-07 | Mālpils (LVA) |
| *S. balticum* | 41118 | 1.1, **1.2**, 2.2, 3.3, 8.1, 9.5 | 2016-08 | Lapland (SWE)* |
| *S. capillifolium* | 41126 | 1.2, 1.3, 1.5, **1.8**, 1.9, 1.49 | 2015-07 | Freiburg, Schauinsland (DEU)* |
| *S. centrale* | 41129 | 1.2, 3.3, 6.4, 7.5 | 2016-08 | Siberia, Surgut Polesye (RUS) |
|  | 41134 | 9.6, **10.2** | 2016-07 | Mālpils (LVA) |
| *S. compactum* | 41137 | 3.1, 4.6, **5.1**, 5.3, 6.1, 6.2 | 2018-06 | Bargerveen (NLD) |
| *S. cuspidatum* | 41146 | 1.1, 1.4, 3.3, 3.4, 5.1, **5.2** | 2016-07 | Gründlenried - Rötseemoos (DEU)* |
| *S. fallax* | 41151 | 2.1, 3.1, 4.1, 4.4, **4.5**, 4.6 | 2017-06 | Sphagnum farming pilot: Rastede (DEU)<br>Origin: De Werribben (NLD) |
| *S. fimbriatum* | 40069 | **1.1**, 2.1 | 2012-06 | Store Mosse (SWE) |
|  | 41154 | 6.1, 6.2, 6.4, 6.5 | 2015-07 | Mālpils (LVA) |
| *S. fuscum* | 41158 | **1.1**, 2.1, 2.2, 2.3, 3.1, 3.2 | 2016-08 | Lapland (SWE)* |
| *S. lindbergii* | 41167 | 2.1, 2.3, 3.1, **3.2**, 3.3 | 2016-07 | Lapland (SWE)* |
| *S. medium/divinum* | 40066 | 3.1 | 2012-07 | Store Mosse (SWE) |
|  | 41169 | 4.1, 4.2, 4.3, 5.1, 5.2 | 2016-08 | Siberia, Yugra (RUS) |
| *S. palustre* | 40068 | 2a, **12a** | 2012-08 | Lychen-Bohmshof (DEU) |
|  | 41175 | 4.2, 4.3, 5.1, 5.2 | 2017-06 | Sphagnum farming pilot: Rastede (DEU)<br>Origin: De Werribben (NLD) |
| *S. papillosum* | 41179 | 1.1, 2.2, 4.3 | 2016-08 | Siberia, Potanay Aapa mire (RUS) |
|  | 41183 | 5.2, **6.1**, 7.1 | 2017-06 | Sphagnum farming pilot: Rastede (DEU)<br>Origin: Ramsloh (DEU) |
| *S. rubellum* | 40067 | **1.1**, 2.1 | 2012-06 | Store Mosse (SWE) |
| *S. russowii* | 41191 | 1.1, 1.2, 3.1, 3.4, 3.5, **4.2** | 2016-08 | Siberia, Chistoye Bog (RUS) |
| *S. squarrosum* | 41193 | 2.1, **5.2**, 5.3, 6.1 | 2016-07 | Gründlenried - Rötseemoos (DEU)* |
|  | 41196 | 7.1 | 2017-05 | Buddenhagener Moor (DEU) |
|  | 41197 | 8.3 | 2017-09 | Steiermark (AUT) |
| *S. subnitens* | 40070 | 1.1 | 2012-06 | Store Mosse (SWE) |
| *S. subfulvum* | 41201 | 4.2, 4.3, 7.2, **7.4**, 8.1, 8.5 | 2016-08 | Lapland (SWE)* |
| *S. warnstorffii* | 41208 | 1.3, 1.4, 2.1, 3.3, **5.2**, 5.4 | 2016-08 | Siberia, Rangetur Floating mire (RUS) |

The six best-growing clones are listed by the number of spore capsule and of the individual clone, respectively. Bold numbers indicate the best-growing clones. The origin of the spore capsule is indicated by date and location of collection. Spore capsules marked with an asterisk (\*) were provided by Michael Lüth, all others by the authors. AUT = Austria, DEU = Germany, LVA = Latvia, NLD = Netherlands, RUS = Russian Federation, SWE = Sweden.

### Cell-cycle arrest, genome sizes and ploidy

The DNA content of the nuclei (ploidy) can affect productivity, at least in animals and seed plants (Dhawan & Lavania, 1996; Chen, 2013; Paterson *et al*., 2012). Usually, *Sphagnum* species have haploid gametophytes and n = 19 chromosomes, while diploid forms with 38 chromosomes exist. Both chromosome numbers exist for populations of some species (Cronberg, 1993). Besides chromosome counting, *Sphagnum* genome sizes were estimated by Feulgen absorbance microscopy (Temsch *et al*., 1998). Flow cytometry (FCM) was applied to determine DNA contents of mosses (Reski *et al*., 1994), including *Sphagnum* (Melosik *et al*., 2005). The major peak of the internal standard *Physcomitrella patens* represents haploid nuclei in the G2-phase of the cell cycle (Schween *et al*., 2003a). It was set at channel 200, whereas the peak at channel 100 represents nuclei in G1. A peak at 400 indicates diploid nuclei in G2 (Schween *et al*., 2003a).

Our FCM analysis revealed only one peak for gametophytic cells of all 19 *Sphagnum* species, but at two different positions: Either one peak occurred around 100, or a peak occurred near 200 (**Fig. 5**). As we analysed fast-growing tissue one might expect that nuclei from one sample were in different phases of the cell cycle and thus would yield two different peaks (G1 and G2) plus intermediary signals for nuclei in the S-phase. Our current findings confirm similar findings of Melosik *et al*. (2005) and suggest that gametophytic cells of all 19 *Sphagnum* species are arrested predominantly either in G1 or G2. A similar cell-cycle arrest occurs in *P. patens* (Reski *et al*., 1994; Schween *et al*., 2003a).

**Fig. 5.**
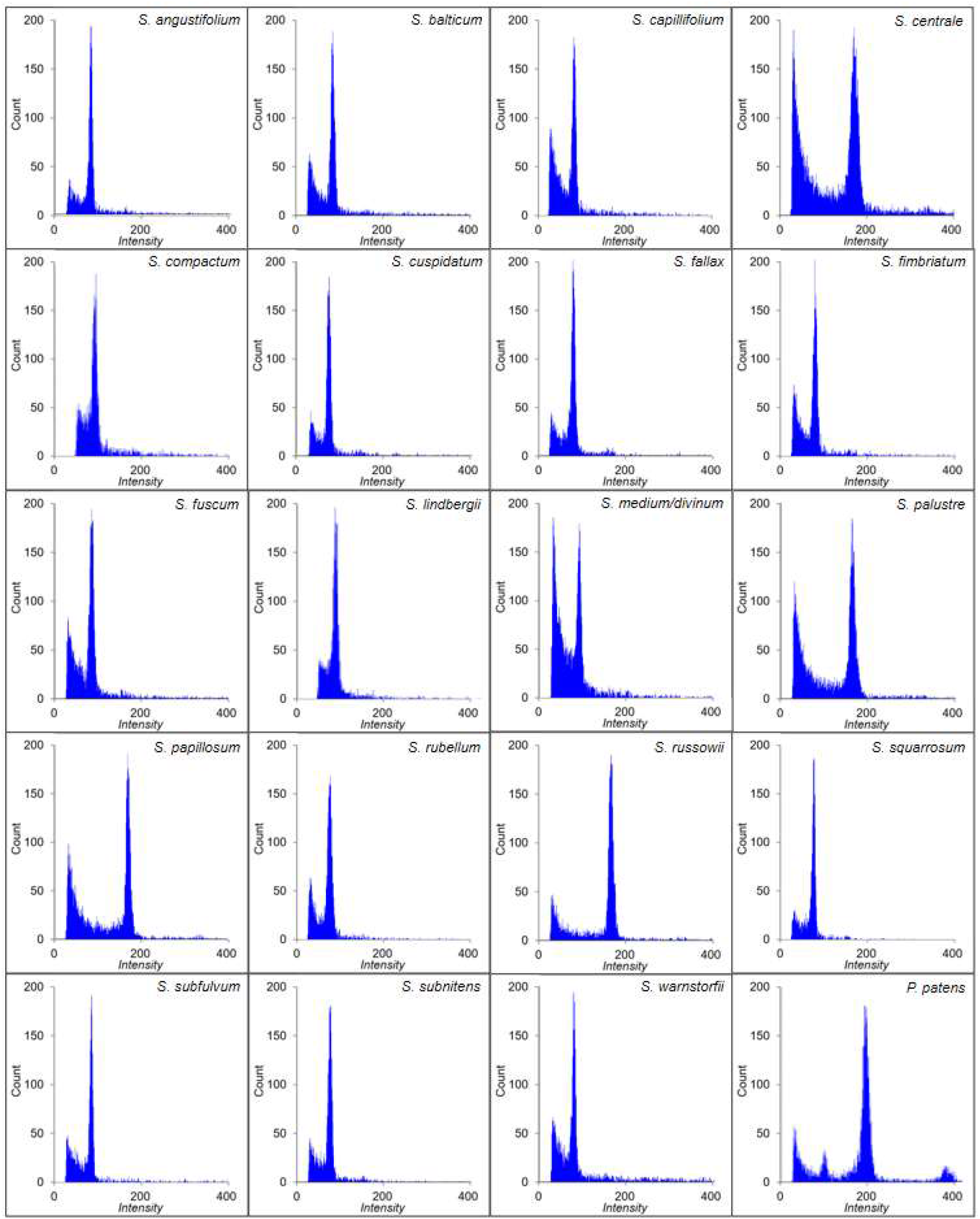
Flow-cytometry (FCM) signals of 19 *Sphagnum* species and *Physcomitrella patens* after axenic cultivation on solid Sphagnum medium for four months. *P. patens* was used as internal standard with the major peak set at channel 200. Channel numbers (x-axis) reflect the relative fluorescence intensity of the stained nuclei.

In such a situation, FCM cannot clarify if a peak corresponds to G1 or to G2. Thus, a peak at 200 could result from either haploid nuclei in G2, or diploid nuclei in G1. Thus, we considered the published genome sizes that are based on sequencing: The basic nuclear DNA content of *P. patens* is 0.53 pg, with an estimated genome size of 518 megabase pairs (Mbp) (Schween *et al*., 2003a), while the latest *P. patens* genome assembly yielded 467.1 Mbp (Lang *et al*., 2018). Based on Feulgen absorbance photometry, haploid *Sphagnum* species have DNA contents between 0.392 pg and 0.506 pg, while diploid species have between 0.814 and 0.952 pg DNA (Temsch *et al*., 1998) with an average ratio of 1:1.92 between DNA content in haploids and diploids (Melosik *et al*., 2005). Currently, there is only one *Sphagnum* genome sequence publicly available. According to this, the *S. fallax* genome comprises approximately 395 Mbp (http://phytozome.jgi.doe.gov/). Assuming that in our FCM analysis peaks at 100 and at 200 represent cells in G1, the estimated genome sizes vary between 370 and 460 Mbp for the peak at 100 and between 840 and 890 Mbp for the peak at 200. Although these are only approximations because DAPI binds to AT-rich DNA sequences (Doležel *et al*., 1992), the values for those *Sphagnum* species characterised by a peak around 100 coincide well with the size of the *S. fallax* genome. We therefore conclude that the gametophytic cells of these species are haploid and predominantly arrested in G1.

Although it is an obvious hypothesis that species with a peak around 200 are diploid and arrested in G1, and not haploid and arrested in G2, we tested this hypothesis by comparison with the literature about haploidy and diploidy in *Sphagnum* and compiled the data in **Table 2**. Our hypothesis is in accordance with data for the 13 haploid species *S. angustifolium, S. balticum, S. capillifolium, S. compactum, S. cuspidatum, S. fallax, S. fuscum, S. lindbergii, S. medium/divinum*, *S. rubellum, S. subnitens, S. subfulvum* and *S. warnstorfii*, as well as the three diploid species *S. centrale, S. palustre* and *S. russowii*. Our *S. fimbriatum* clones, which derive from sporophytes collected in Sweden and Latvia are haploid. *S. fimbriatum* was reported to be haploid in the USA (Bryan, 1955), Finland (Sorsa, 1955, 1956), Canada (Maass & Harvey, 1973) and Austria (Temsch *et al*., 1998), whereas diploid specimens were reported for the UK (Smith & Newton, 1968). Our *S. papillosum* clones established from sporophytes collected in Russia and Germany are diploid, which is in agreement with material from the UK (Smith & Newton, 1968) and Austria (Temsch *et al*., 1998), whereas haploid specimens were reported from Canada (Maass & Harvey, 1973). Our *S. squarrosum* clones from Germany and Austria are haploid, like material from Austria (Temsch et al., 1998) and Canada (Maass & Harvey, 1973), whereas diploid specimens were reported from Finland (Sorsa, 1955, 1956).

Taken together, we conclude that the gametophytic cells of 19 *Sphagnum* species are predominantly arrested in G1, at least under our conditions. This contrasts with the G2-arrest of *Physcomitrella patens* protonemal cells. One prominent feature of *P. patens* is the very high efficiency of homologous recombination (HR) in these cells. This feature facilitates precise gene targeting (GT) and thus genome engineering with outstanding efficiency (Schaefer & Zryd, 1997; Strepp *et al*., 1998; Hohe *et al*., 2004). Although it is not yet fully resolved why *P. patens* has such an outstandingly high HR-efficiency, two hypotheses were put forward early on: either haploidy or the G2-arrest is a prerequisite (Schaefer & Zryd, 1997; Reski, 1998). These hypotheses can be tested with the collection described here: If haploidy is sufficient, haploid but not diploid *Sphagnum* species should be amenable to efficient GT. If G2-arrest is a prerequisite, none of the species described here is amenable to GT. To our knowledge, no genetic transformation of any *Sphagnum* species was hitherto reported. Because protoplasts derived from protonemal cells are the preferred target for genetic transformation in *P. patens*, the report about protonema-induction in *S. squarrosum* (Zhao *et al*., 2019) paves the way for such experiments in the future.

**Table 2.** The ploidy level of 19 *Sphagnum* species measured by FCM in comparison with the literature

| Species | This study<br>FCM | Level of ploidy |  |  |  |  |  |
| --- | --- | --- | --- | --- | --- | --- | --- |
|  |  | Bryan,<br>1955 | Sorsa,<br>1955 | Sorsa,<br>1956 | Smith and<br>Newton, 1968 | Maass and<br>Harvey, 1973 | Temsch et<br>al., 1998 |
| *S. angustifolium* | n |  |  |  |  |  | n |
| *S. balticum* | n |  | n | n |  |  |  |
| *S. capillifolium* | n |  |  |  |  |  | n |
| *S. centrale* | 2n |  |  |  |  | 2n | 2n |
| *S. compactum* | n | n | n | n |  |  | n |
| *S. cuspidatum* | n | n | n | n | n | n | n |
| *S. fallax* | n |  |  |  |  |  | n |
| *S. fimbriatum* | n | n | n | n | 2n | n | n |
| *S. fuscum* | n |  | n | n |  | n | n |
| *S. lindbergii* | n |  |  | n |  | n |  |
| *S. magellanicum* |  | n |  | 2n |  | n | n |
| *S. medium/divinum* | n |  |  |  |  |  |  |
| *S. palustre* | 2n | 2n |  |  | 2n | 2n | 2n |
| *S. papillosum* | 2n |  |  |  | 2n | n | 2n |
| *S. rubellum* | n |  |  |  | n | n | n |
| *S. russowii* | 2n |  |  |  |  |  | 2n |
| *S. squarrosum* | n |  | 2n | 2n |  | n | n |
| *S. subnitens* | n |  | n |  | n |  | n |
| *S. subfulvum* | n |  |  |  |  | n |  |
| *S. warnstorffii* | n |  | n | n |  |  | n |

Literature data are based on counting chromosome numbers (Bryan, 1955; Sorsa, 1955, 1956; Smith & Newton, 1968; Maass & Harvey, 1973) or on genome size determination with Feulgen absorbance photometry and a scanning cytophotometer (Temsch *et al.*, 1998). n=haploid, 2n=diploid.

### Growth in suspension

Productivity of *Sphagnum* in their natural habitats varies among species with a biomass production of up to 1450 g m^−2^ y^−1^ with an average of 260 g m^−2^ y^−1^, depending on phylogeny and microhabitat preferences (Gunnarsson, 2005). To compare productivity without the influence of water level or nutrient supply, we tested cultivation in suspensions under standardized conditions to identify the best-growing clone of each of the 19 species (**Fig. 6**). To compare the growth behaviour of the species, the inocula have to be normalized. Due to the variation of the capitula sizes of *Sphagnum* species (**Fig. 2**), the inoculation material had to be adjusted. *In-vitro* cultures facilitate the reproduction due to vegetative growth, because peat mosses regenerate from several parts of the shoot like capitula, fascicles, branches and stems, but not from leaves (Poschlod & Pfadenhauer, 1989).

**Fig. 6.**
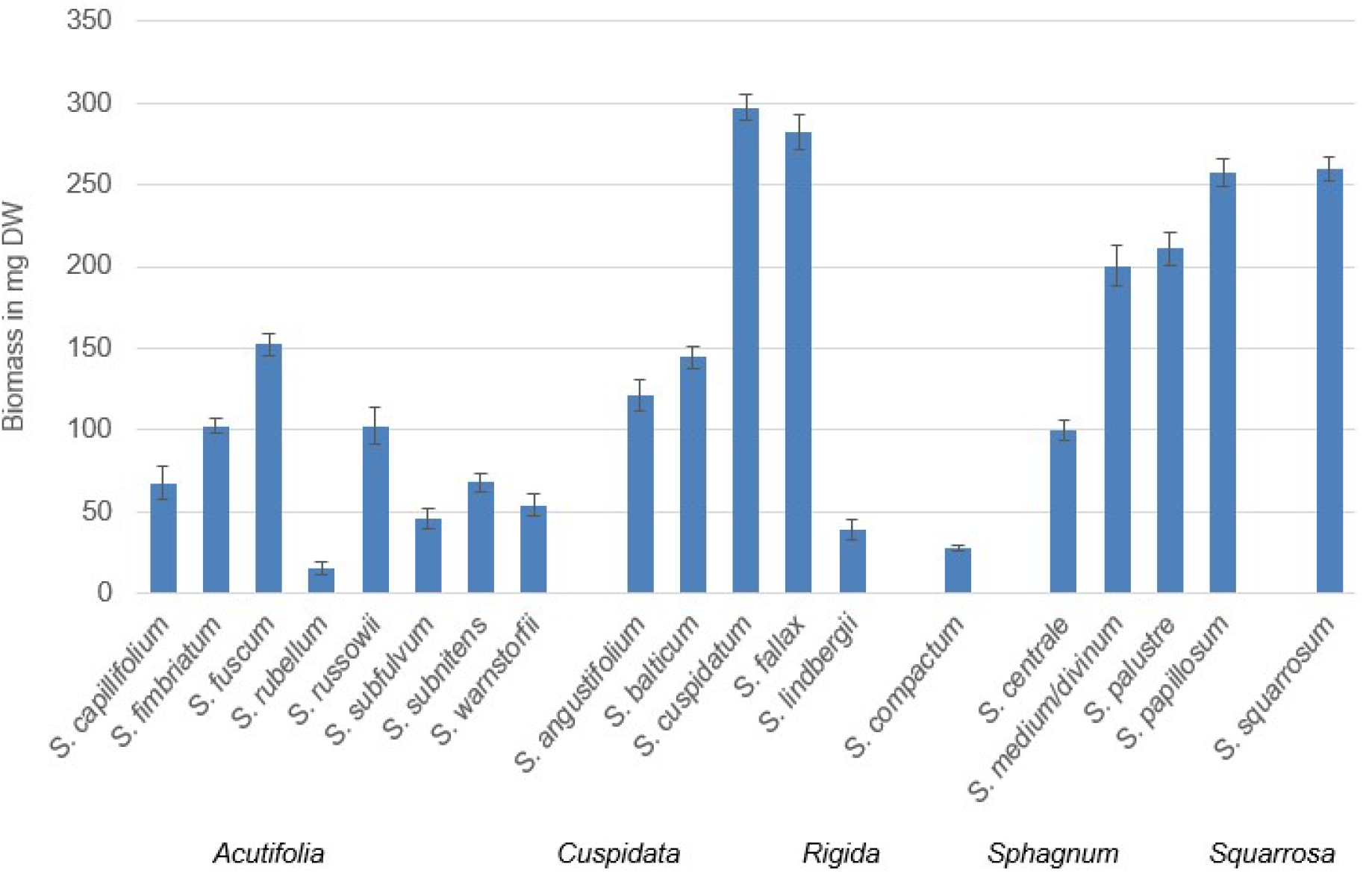
Biomass increase of 19 *Sphagnum* species sorted by sections after cultivating 50 mg FW (≈3.6 mg DW) of gametophores in flasks containing 35 ml Sphagnum medium for six weeks. The y-axis shows the biomass in mg dry weight, the x-axis shows *Sphagnum* species. Data represent mean values with standard deviations of three biological replicates (n=3).

We disrupted gametophores with forceps and filled flasks with 50 mg fresh weight (FW; ≈3.6 mg DW cf. Beike *et al*., 2015) and 35 ml Sphagnum medium. The nutrient composition was established towards optimized biomass production of *S. palustre*, but it was not proven for the other established axenic *in-vitro* cultures of *S. fimbiratum, S. magellanicum, S. rubellum* and *S. subnitens* (Beike *et al*., 2015). Under our conditions, biomass increase ranged from 4-fold (*S. rubellum*) up to 80-fold (*S. cuspidatum*) in six weeks (**Fig. 6**).

The Sphagnum medium was suitable for axenic *in-vitro* cultivation of many *Sphagnum* species, including *S. cuspidatum* (yielding the largest biomass gain), *S. fallax, S. papillosum* and *S. squarrosum* (**Fig. 6**). The medium seems suboptimal for e.g. *Sphagnum subnitens*, the species with the highest productivities out of 31 peat-moss species from Sphagnum-dominated wetlands (Gunnarsson, 2005). However, in our study *S. subnitens* had only a weak performance. Gaudig *et al*. (2020) reported that *S. papillosum, S. palustre, S. fimbriatum* and *S. fallax* grow well under nutrient-rich conditions with optimal water supply in a glasshouse experiment. In this study, *S. fallax* had the highest and *S. papillosum* the lowest productivity at high water level. The good productivity of *S. fimbriatum* in the glasshouse experiment contrasts with its comparably low productivity in suspension in our study. In the field, biomass increase is generally largest in pools, less on lawns and least on hummocks (Clymo, 1970), with species growing in ombrotrophic carpets and lawns like *S. balticum, S. cuspidatum, S. magellanicum* and *S. rubellum* having higher productivities than hummock species like *S. fuscum* (Gunnarsson, 2005). This corresponds with several studies, which have shown that the growth rate of most *Sphagnum* species is highest at water tables just below the capitula, independent of the species (e.g. Gaudig *et al*., 2020). Interestingly, we recorded higher growth rates for *S. papillosum* than for *S. palustre* although the opposite is described for natural habitats (Gunnarsson, 2005; Krebs *et al*., 2016). Surprisingly, axenic *in-vitro* cultivation enhances the productivity of *S. fuscum*, but impairs the growth of *S. rubellum*.

We could not correlate productivity and taxonomical sections as the four most productive species in our study belong to three different sections (*Cuspidata, Sphagnum, Squarrosa*). Species from the sections *Acutifolia* and *Rigida* had a lower average productivity. In natural habitats, species of the section *Cuspidata* are more productive than species of the sections *Acutifolia* and *Sphagnum* (Gunnarsson, 2005). To clarify the differences in productivity on a genetic level between the taxonomical sections, a larger number of species from one section should be examined in future. Our inability to detect a correlation between taxonomical sections and biomass gain may reflect the situation in the field. Piatkowski & Shaw (2019) did not detect an influence of phylogeny on the majority of traits in their study on 15 *Sphagnum* species and suggest that the environmental context can obscure the phylogenetic signal. Our novel method creates an artificial but standardized environment for 19 *Sphagnum* species. It facilitates research to gain deeper insights into the ecology of peat mosses as single parameters like nutrients and light conditions can be changed and tested. Recently, Küttim et al. (2020) reported on biomass increases of *Sphagnum* species during boreal winters. Consequently, future studies should also take climate gradients into account.

Another genetic property that influences productivity is ploidy (Otto & Whitton, 2000), as described for many agricultural crops (Henry & Nevo, 2014). In our current study, however, we could not detect a correlation between ploidy and productivity, because the diploid *S. palustre* and *S. papillosum* were among the six best-growing species, but the haploid *S. cuspidatum* and *S. fallax* were most productive. The other diploid species *S. centrale* and *S. russowii* yielded an average increase. Because the medium composition may have affected our results, future studies of haploid and diploid populations of the same species using individually optimized media may provide better insights into a correlation between ploidy and yield. Alternatively, genome duplication may have other benefits than pure biomass increase in *Sphagnum*, or even mosses in general.

### Summary results of axenic cultivation of 19 *Sphagnum* species

#### *Sphagnum angustifolium* (Section *Cuspidata*)

Six clones were obtained from two sporophytes collected in Latvia. The productivity of clones 2.1, 2.2, 2.3, 2.4, 2.5, and 7.1 in suspension ranged from 34.4±6 mg DW (clone 2.2) to 226.6±7.3 mg DW (clone 2.4). Clone 2.4 yielded significantly more biomass compared to the other five (Figure S1). All clones are haploid, which is consistent with Temsch *et al*. (1998).

#### *Sphagnum balticum* (Section *Cuspidata*)

16 clones were obtained from six sporophytes collected in Sweden. Clones 1.1, 1.2, 2.2, 3.3, 8.1 and 9.5 were the six best-growing clones on solid medium. Growth in suspension resulted in the same productivity for five clones, while clone 1.2 grew more than twice as fast and yielded significantly more biomass (Figure S2). All clones are haploid, which is consistent with Sorsa (1955, 1956).

#### *Sphagnum capillifolium* (Section *Acutifolia*)

11 clones were obtained from one sporophyte collected in Germany. Clones 1.2, 1.3, 1.5, 1.8, 1.9 and 1.49 were the six best-growing clones on solid medium. Growth in suspension ranged from 6.9±0.8 mg DW (clone 1.2) to 45.7±8.2 mg DW (clone 1.8). Clone 1.8 yielded significantly more biomass compared to the other five (Figure S3). All clones are haploid, which is consistent with Temsch *et al*. (1998).

#### *Sphagnum centrale* (Section *Sphagnum*)

14 clones were obtained from seven sporophytes collected in Russia (clones 1-7) and Latvia (9-10). Clones 1.2, 3.3, 6.4, 7.5, 9.6 and 10.2 were the six best-growing clones on solid medium. Growth in suspension yielded an average biomass of 119.3±11.4 mg DW, while clone 10.2 was significantly more productive than the other five (Figure S4). All clones are diploid, which is consistent with the literature (Temsch *et al*., 1998; Maass & Harvey, 1973).

#### *Sphagnum compactum* (Section *Rigida*)

12 clones were obtained from four sporophytes collected in the Netherlands. Clones 3.1, 4.6, 5.1, 5.3, 6.1 and 6.2 were the six best-growing clones on solid medium. Growth in suspension ranged from 4.0±1.8 mg DW (clone 6.2) to 33.2±7.7 mg DW (clone 5.1). The best-growing clone 5.1 yielded significantly more biomass than clones 3.1, 5.3, 6.1 and 6.2, but not significantly more than clone 4.6 (Figure S5). All clones are haploid, which is consistent with the literature (Bryan, 1955; Sorsa, 1955, 1956; Temsch *et al*., 1998).

#### *Sphagnum cuspidatum* (Section *Cuspidata*)

15 clones were obtained from of five sporophytes collected in Germany. Clones 1.1, 1.4, 3.3, 3.4, 5.1 and 5.2 were the six best-growing clones on solid medium. Growth in suspension ranged from 58.4±4.6 mg DW (clone 3.3) to over 150 mg DW (clones 1.4, 5.2). The best-growing clone 5.2 produced significantly more biomass than clones 1.1, 3.3, 3.4 and 5.1, but not significantly more than clone 1.4 (Figure S6). All clones are haploid, which is consistent with the literature (Bryan, 1955; Sorsa, 1955, 1956; Smith & Newton, 1968; Maass & Harvey, 1973; Temsch *et al*., 1998).

#### *Sphagnum fallax* (Section *Cuspidata*)

10 clones were obtained from three sporophytes collected on a Sphagnum farming pilot field in Germany, where in 2011 fragments of a mixture of *Sphagnum* species collected in the Netherlands were spread out on former bog grassland (Gaudig *et al*., 2014). Clones 2.1, 3.1, 4.1, 4.4, 4.5 and 4.6 were the six best-growing clones on solid medium. Growth in suspension ranged from 23.5±5.1 mg DW (clone 2.1) to over twice that amount (clone 4.5). Biomass increase of the best-growing clone 4.5 was not significantly higher than that of clones 3.1, 4.1, 4.4 and 4.6 (Figure S7). All clones are haploid, which is consistent with Temsch *et al*. (1998).

#### *Sphagnum fimbriatum* (Section *Acutifolia*)

Six clones were obtained from three sporophytes from Sweden (1-2) and Latvia (6). The clones 1.1 and 2.1 were established by Beike *et al*. (2015). Growth in suspension of clones 1.1, 2.1, 6.1, 6.2, 6.4 and 6.5 ranged from 23.9±3.8 mg DW (clone 2.1) to 60.2±3.9 mg DW (clone 1.1). The best-growing clone 1.1 was significantly more productive than clones 2.1, 6.1 and 6.4, but not significantly more than clones 6.2 and 6.5 (Figure S8). All clones are haploid, which is consistent with Bryan (1955), Sorsa (1955, 1956), Maass & Harvey (1973) and Temsch *et al*. (1998), but in contrast to Smith & Newton (1968).

#### *Sphagnum fuscum* (Section *Acutifolia*)

Eight clones were obtained from three sporophytes collected in Sweden. Clones 1.1, 2.1, 2.2, 2.3, 3.1 and 3.2 were the six best-growing clones on solid medium. Growth in suspension yielded about the same productivity for five clones, whilst clone 1.1 was significantly twice as productive (Figure S9). All clones are haploid, which is consistent with the literature (Sorsa, 1955, 1956; Maass & Harvey, 1973; Temsch *et al*., 1998).

#### *Sphagnum lindbergii* (Section *Cuspidata*)

Five clones were obtained from two sporophytes collected in Germany. Growth in suspension of clones 2.1, 2.3, 3.1, 3.2 and 3.3 yielded from 21.9±1.4 mg DW (clone 3.3) to over 40 mg DW (clones 2.1, 3.2). The best-growing clone 2.1 was significantly more productive than clones 2.3 and 3.3, but not significantly more than clones 3.1 and 3.2 (Figure S10). All clones are haploid, which is consistent with the literature (Sorsa, 1956; Maass & Harvey, 1973).

#### *Sphagnum medium/divinum* (Section *Sphagnum*)

Eight clones were obtained from four sporophytes collected in Sweden (1, 3) and Russia (4-5). Clone 3.1 was established as *S. magellanicum* by Beike *et al*. (2015). Clones 3.1, 4.1, 4.2, 4.3, 5.1 and 5.2 were the six best-growing clones on solid medium. Growth in suspension yielded from 28.4±7.1 mg DW (clone 4.3) to twice that amount (clones 3.1 and 4.1). The best-growing clone 3.1 was more productive than clones 4.2, 4.3, 5.1 and 5.2, but not significantly more than clone 4.1 (Figure S11). All clones are haploid, which is consistent with Bryan (1955), Maass & Harvey (973) and Temsch *et al*. (1998) but contrasts Sorsa (1956), who reported diploid specimens. All these reports named the species *S. magellanicum*.

#### *Sphagnum palustre* (Section *Sphagnum*)

Nine clones were obtained from three sporophytes collected from different locations in Germany. Clones 2a and 12a germinated out of one spore capsule established by Beike *et al*. (2015) under cultivation conditions optimized for clone 12a. Clones 2a, 12a, 4.2, 4.3, 5.1 and 5.2 were the six best-growing clones on solid medium. Suspension culture yielded an average biomass of 155.5±6.5 mg DW, with clone 12a being most productive, but not significantly more than clones 4.2, 4.3, 5.1 and 5.2 (Figure S12). All clones are diploid, which is consistent with the literature (Bryan, 1955; Smith & Newton, 1968; Maass & Harvey, 1973; Temsch *et al*., 1998).

#### *Sphagnum papillosum* (Section *Sphagnum*)

12 clones were obtained from six sporophytes collected in Russia (1-4) and on a Sphagnum farming field in Rastede, Germany (5-7), established in 2011 from a mixture of *Sphagnum* species collected in Ramsloh, Germany (Gaudig *et al*., 2014). Clones 1.1, 2.2, 4.3, 5.2, 6.1 and 7.1 were the six best-growing clones on solid medium. Growth in suspension yielded from 26.1±5.8 mg DW (clone 7.1) to 57.0±10.1 mg DW (clone 6.1). The best-growing clone 6.1 was significantly more productive than clones 2.2, 4.3, 5.2, and 7.1, but not significantly more than clone 1.1 (Figure S13). All clones are diploid, which is consistent with Smith & Newton (1968) and Temsch *et al*. (1998), whereas Maass & Harvey (1973) reported haploid specimens.

#### *Sphagnum rubellum* (Section *Acutifolia*)

Two clones were obtained from two sporophytes collected in Sweden. Clone 1 was selected as best-growing clone, it was established by Beike *et al*. (2015). Both clones are haploid, which is consistent with the literature (Smith & Newton, 1968; Maass & Harvey, 1973; Temsch *et al*., 1998).

#### *Sphagnum russowii* (Section *Acutifolia*)

12 clones were obtained from three sporophytes collected in Russia. Clones 1.1, 1.2, 3.1, 3.4, 3.5 and 4.2 were the six best-growing clones on solid medium. Growth in suspension ranged from 27.9±1.1 mg DW (clone 3.5) to 76.8±12.1 mg DW (clone 4.2). Clone 4.2 was significantly more productive than the other five (Figure S14). All clones are diploid, which is consistent with Temsch *et al*. (1998).

#### *Sphagnum squarrosum* (Section *Squarrosa*)

16 clones were obtained from six sporophytes collected in Germany and Austria. Clones 2.1, 5.2, 5.3, 6.1, 7.1 and 8.3 were the six best-growing clones on solid medium. Growth in suspension yielded an average biomass of 142.9±18.1 mg DW. The best-growing clone 5.2 yielded significantly more biomass than clones 2.1, 5.3, 6.1 and 8.3, but not significantly more than clone 7.1 (Figure S15). All clones are haploid, which is consistent with Maass & Harvey (1973) and Temsch *et al*. (1998), but contradicts Sorsa (1955, 1956).

#### *Sphagnum subfulvum* (Section *Acutifolia*)

16 clones were obtained from six sporophytes collected in Sweden. Clones 4.2, 4.3, 7.2, 7.4, 8.1 and 8.5 were the six best-growing clones on solid medium. Growth in suspension yielded an average biomass of 13.0±4.2 mg DW, with clone 7.4 being the best-growing clone (Figure S16). All clones are haploid, which is consistent with Maass & Harvey (1973).

#### *Sphagnum subnitens* (Section *Acutifolia*)

One clone was obtained from a sporophyte collected in Sweden and was established by Beike et al. (2015). It is haploid, which is consistent with the literature (Sorsa, 1955, 1956; Smith & Newton, 1968; Temsch *et al*., 1998).

#### *Sphagnum warnstorfii* (Section *Acutifolia*)

14 clones were obtained from four sporophytes collected in Russia. Clones 1.3, 1.4, 2.1, 3.3, 5.2 and 5.4 were the six best-growing clones on solid medium. Growth in suspension ranged from 16.1±4.9 mg DW (clone 3.5) to 34.4±5.4 mg DW (clone 4.2). The best-growing clone 5.2 yielded significantly more biomass than clones 1.3 and 2.1, but not significantly more than clones 1.4, 3.3 and 5.4 (Figure S17). All clones are haploid, which is consistent with the literature (Sorsa, 1955, 1956; Temsch *et al*., 1998).

## Conclusion

Apart from *P. patens* as an established model organism, the development of other model mosses is inevitable for ecological and evolutionary genomics, as well as clarifying open questions such as the high HR efficiency of *P. patens*. Due to the large number of peat-moss species and their clear patterns of niche differentiation, *Sphagnum* provides an exceptional complement (Shaw *et al*., 2016). Our peat-moss collection creates a resource for the increasing interest in *Sphagnum* research to establish new plant model systems. Moreover, the large-scale implementation for diverse applications can rely on the axenic *in-vitro* cultivation of *Sphagnum* as fast growing high-quality founder material. Scaling up this cultivation method will facilitate a low cost production process. Especially Sphagnum farming will benefit, as the lack of *Sphagnum* diaspores is one of the biggest problems and their purchase is the biggest cost factor for establishing Sphagnum farming sites (Wichmann *et al*., 2017, 2020).

## Acknowledgements

This work was supported by the German Federal Ministry of Food and Agriculture (BMEL) (MOOSzucht, No. 22007216). Additional support came from the German Research Foundation (DFG) under Germany’s Excellence Strategy (CIBSS – EXC-2189 – Project ID 390939984; *liv*MatS – EXC-2193/1 – 390951807). We gratefully acknowledge Anja Kuberski for expert technical assistance, Michael Lüth for providing sporophytes, Greta Gaudig for discussion and Anne Katrin Prowse for proofreading of the manuscript.

## Authors contribution

M.A.H., V.M.L., E.L.D. and R.R. planned and designed the research. M.A.H. performed experiments and analysed data. Ma.K., Mi.K. and A.P. collected sporophytes. M.A.H., E.L.D. and R.R. wrote the manuscript. H.J. revised this paper. All authors discussed data and approved the final version of the manuscript.

---

**Fig. S1.**
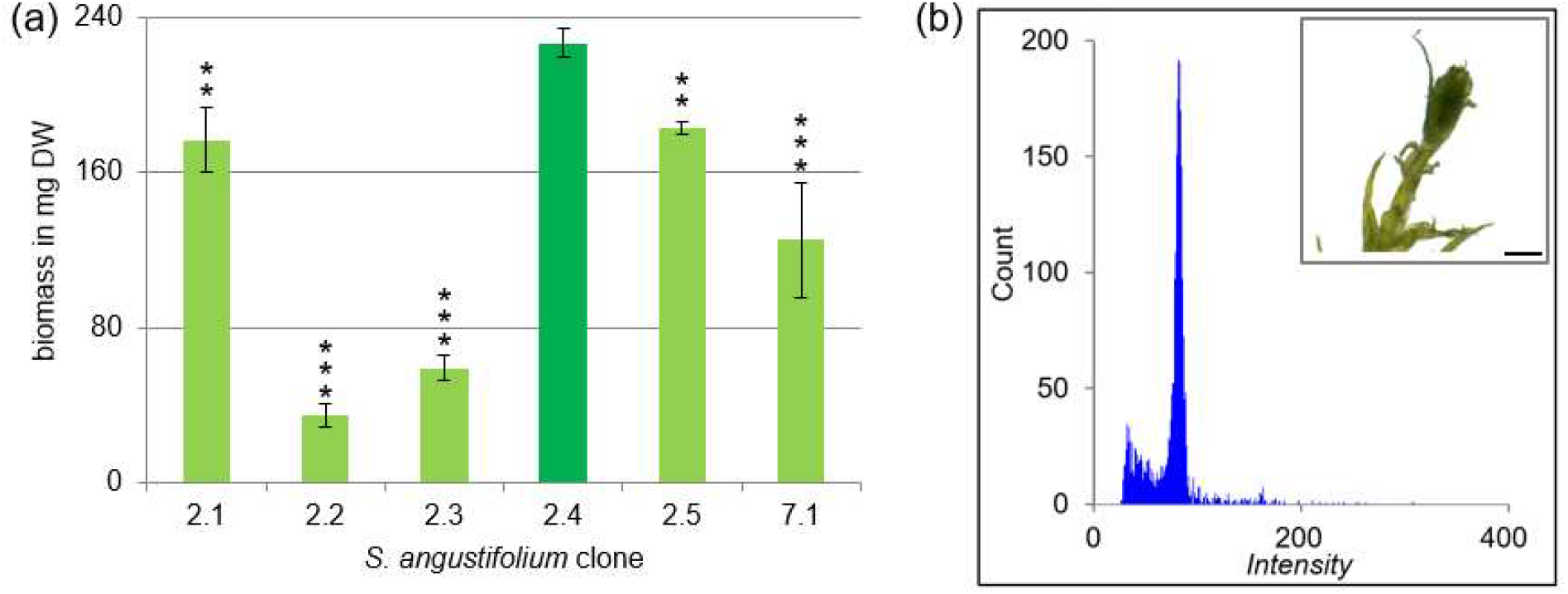
Determination of the best-growing clone of *S. angustifolium*. The growth of the six clones (2.1, 2.2, 2.3, 2.4, 2.5 and 7.1) was determined in suspension (a) by measuring the dry weight after cultivation of three capitula in flasks containing 50 ml Sphagnum media for six weeks. The y-axis shows the biomass in mg dry weight, the x-axis shows the clone. Data represents mean values with standard deviations of 3 biological replicates (ANOVA p<0.0001). Clone 2.4 yielded significantly more biomass compared to clones 2.1**, 2.2***, 2.3***, 2.5** and 7.1***. Asterisks represent results of student t-test performed in comparison to clone 2.4. * = p < 0.05, ** = p < 0.01, *** = p < 0.001. The ploidy of the best-growing clone *S. angustifolium* 2.4 was determined by flow cytometry (FCM). (b) Histograms of gametophore samples measured via FCM after cultivation on solid Sphagnum medium for four months and a picture of *S. angustifolium* gametophore after four weeks (scale bar = 1 mm). The channel numbers corresponding to the relative fluorescence intensities of the analysed particles is shown on the x-axis, the number of counted events is shown on the y-axis.

**Fig. S2.**
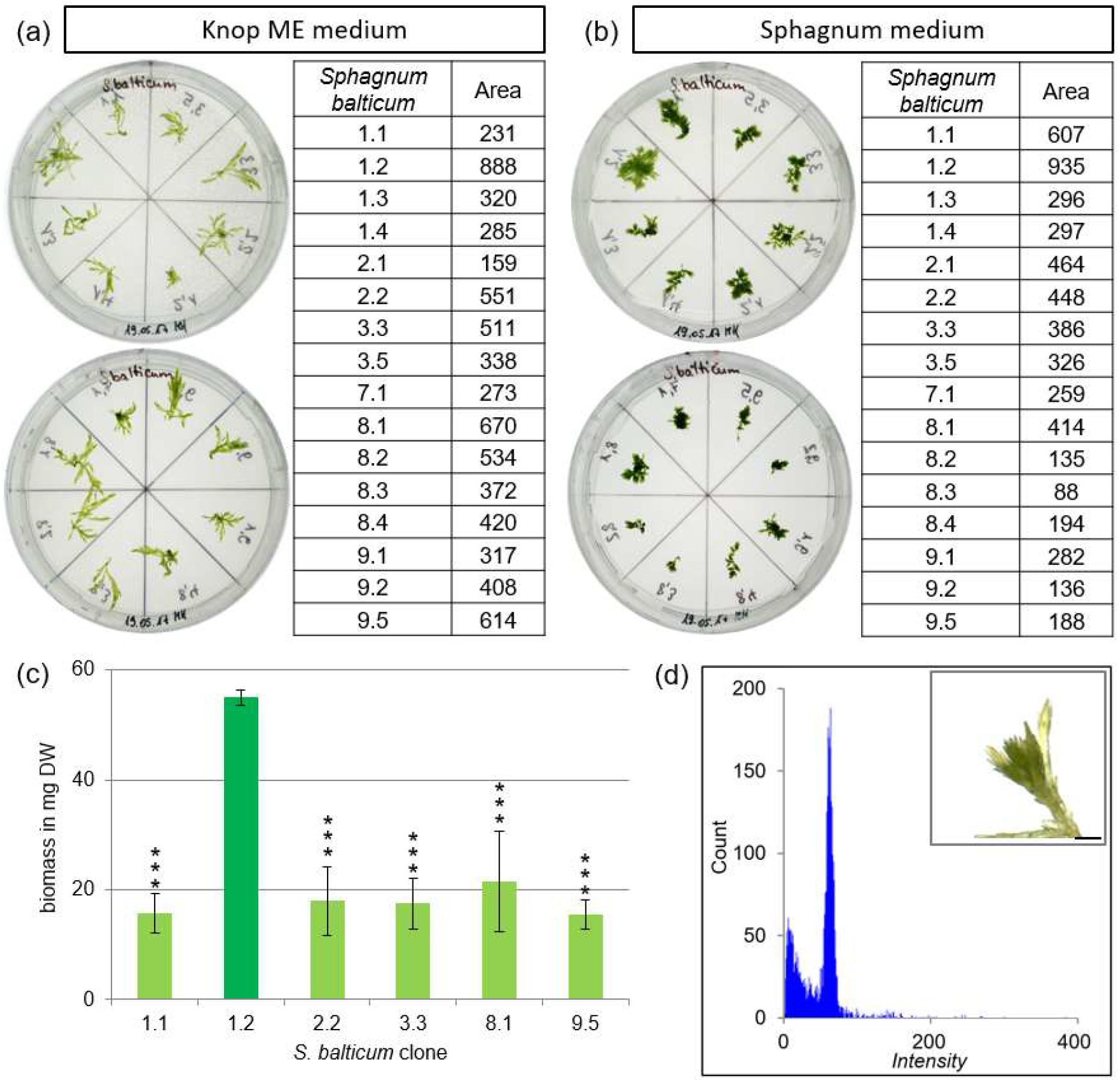
Determination of the best-growing clone of *S. balticum*. Growth determination of 16 *S. balticum* clones on (a) solid Knop ME and (b) solid Sphagnum medium after four weeks of cultivation. The following clones were arranged counterclockwise on the Petri dish: 1.1, 1.2, 1.3, 1.4, 2.1, 2.2, 3.3, 3.5 on upper and 7.1, 8.1, 8.2, 8.3, 8.4, 9.1, 9.2, 9.5 on lower. Gametophores were cultivated for four weeks. The size of the gametophores was measured on the basis of binary pictures using ImageJ and shown in the table next to it. The six best-growing clones (1.1, 1.2, 2.2, 3.3, 8.1 and 9.5) were selected and the growth was determined in suspension (c) by measuring the dry weight after cultivation of three capitula in flasks containing 50 ml Sphagnum media for six weeks. The y-axis shows the biomass in mg dry weight, the x-axis shows the clone. Data represents mean values with standard deviations of 3 biological replicates, except for clone 9.5 (2 replicates) (ANOVA p<0.0001). Clone 1.2 yielded significant more biomass compared to the clones 1.1***, 2.2***, 3.3***, 8.1*** and 9.5***. Asterisks represent results of student t-test performed in comparison to clone 1.2. * = p < 0.05, ** = p < 0.01, *** = p < 0.001. The ploidy of the best-growing clone *S. balticum* 1.2 was determined by flow cytometry (FCM). (d) Histograms of gametophore samples measured via FCM after cultivation on solid Sphagnum medium for four months and a picture of *S. balticum* gametophore after four weeks (scale bar = 1 mm). The channel numbers corresponding to the relative fluorescence intensities of the analysed particles is shown on the x-axis, the number of counted events is shown on the y-axis.

**Fig. S3.**
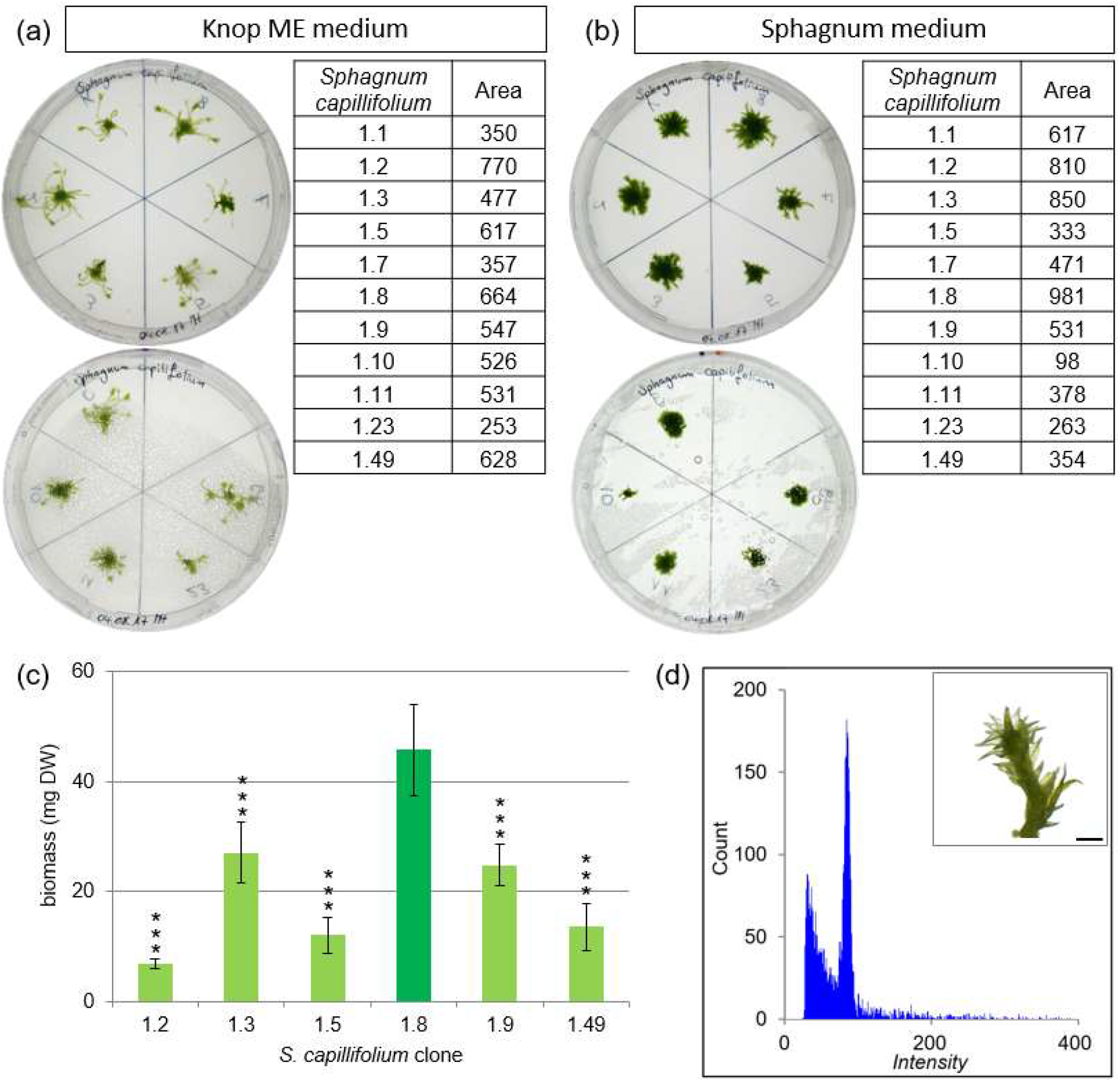
Determination of the best-growing clone of *S. capillifolium*. Growth determination of 11 *S. capillifolium* clones on (a) solid Knop ME and (b) solid Sphagnum medium after four weeks of cultivation. The following clones were arranged counterclockwise on the Petri dish: 1.1, 1.2, 1.3, 1.5, 1.7, 1.8 on upper and 1.9, 1.10, 1.11, 1.23, 1.49 on lower. Gametophores were cultivated for four weeks. The size of the gametophores was measured on the basis of binary pictures using ImageJ and shown in the table next to it. The six best-growing clones (1.2, 1.3, 1.5, 1.8, 1.9 and 1.49) were selected and the growth was determined in suspension (c) by measuring the dry weight after cultivation of three capitula in flasks containing 50 ml Sphagnum media for six weeks. The y-axis shows the biomass in mg dry weight, the x-axis shows the clone. Data represents mean values with standard deviations of 3 biological replicates (ANOVA p<0.0001). Clone 1.8 yielded significant more biomass compared to the clones 1.2***, 1.3***, 1.5***, 1.9*** and 1.49***. Asterisks represent results of student t-test performed in comparison to clone 1.8. * = p < 0.05, ** = p < 0.01, *** = p < 0.001. The ploidy of the best-growing clone *S. capillifolium* 1.8 was determined by flow cytometry (FCM). (d) Histograms of gametophore samples measured via FCM after cultivation on solid Sphagnum medium for four months and a picture of *S. capillifolium* gametophore after four weeks (scale bar = 1 mm). The channel numbers corresponding to the relative fluorescence intensities of the analysed particles is shown on the x-axis, the number of counted events is shown on the y-axis.

**Fig. S4.**
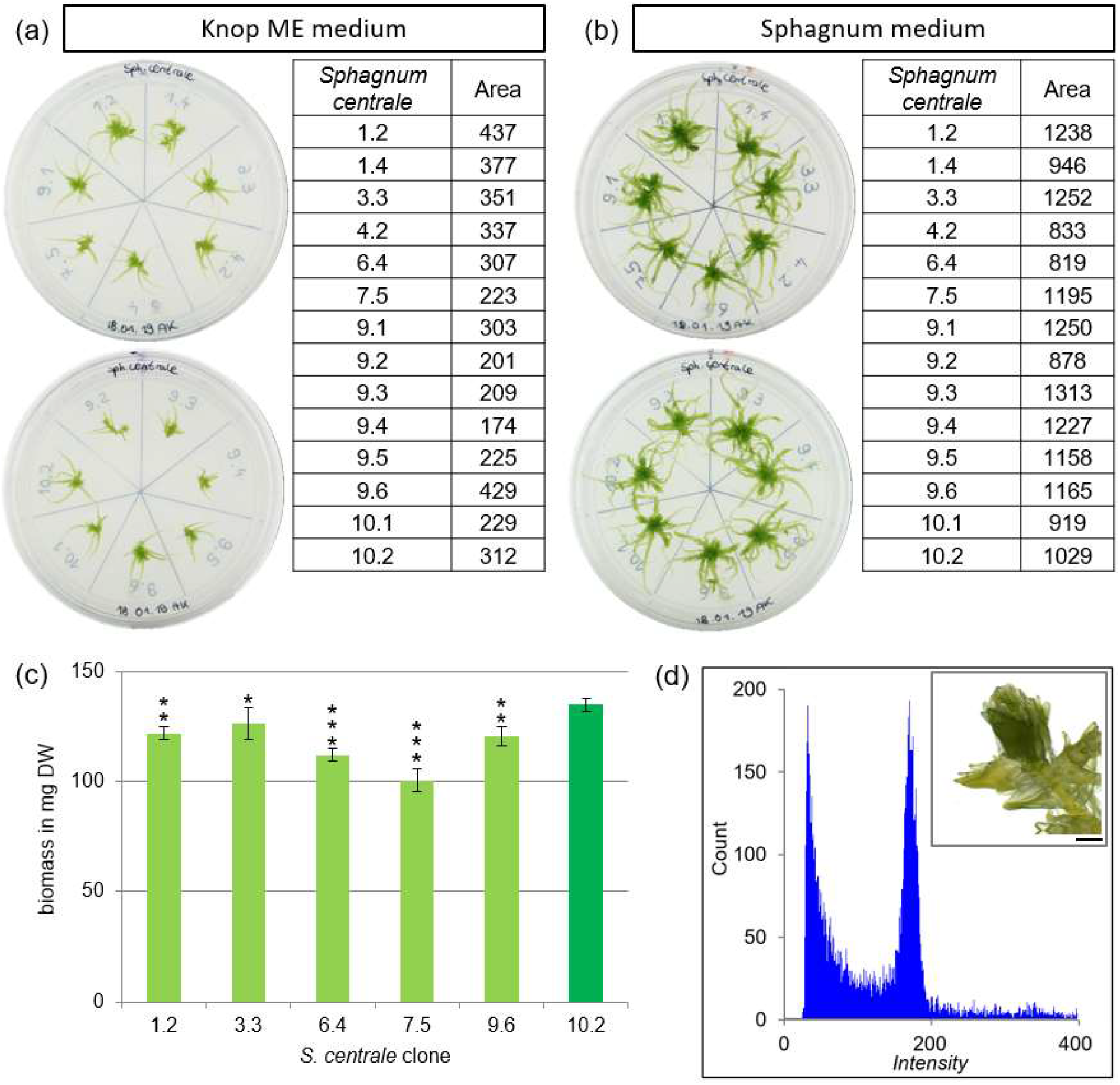
Determination of the best-growing clone of *S. centrale*. Growth determination of 14 *S. centrale* clones on (a) solid Knop ME and (b) solid Sphagnum medium after four weeks of cultivation. The following clones were arranged clockwise on the Petri dish: 1.2, 1.4, 3.3, 4.2, 6.4, 7.5, 9.1 on upper and 9.2, 9.3, 9.4, 9.5, 9.6, 10.1, 10.2 on lower. Gametophores were cultivated for four weeks. The size of the gametophores was measured on the basis of binary pictures using ImageJ and shown in the table next to it. The six best-growing clones (1.2, 3.3, 6.4, 7.5, 9.6 and 10.2) were selected and the growth was determined in suspension (c) by measuring the dry weight after cultivation of three capitula in flasks containing 50 ml Sphagnum media for six weeks. The y-axis shows the biomass in mg dry weight, the x-axis shows the clone. Data represents mean values with standard deviations of 3 biological replicates (ANOVA p<0.0001). Clone 10.2 yielded significant more biomass compared to the clones 1.2**, 3.3*, 6.4***, 7.5*** and 9.6**. Asterisks represent results of student t-test performed in comparison to clone 10.2. * = p < 0.05, ** = p < 0.01, *** = p < 0.001. The ploidy of the best-growing clone *S. centrale* 10.2 was determined by flow cytometry (FCM). (d) Histograms of gametophore samples measured via FCM after cultivation on solid Sphagnum medium for four months and a picture of *S. centrale* gametophore after four weeks (scale bar = 1 mm). The channel numbers corresponding to the relative fluorescence intensities of the analysed particles is shown on the x-axis, the number of counted events is shown on the y-axis.

**Fig. S5.**
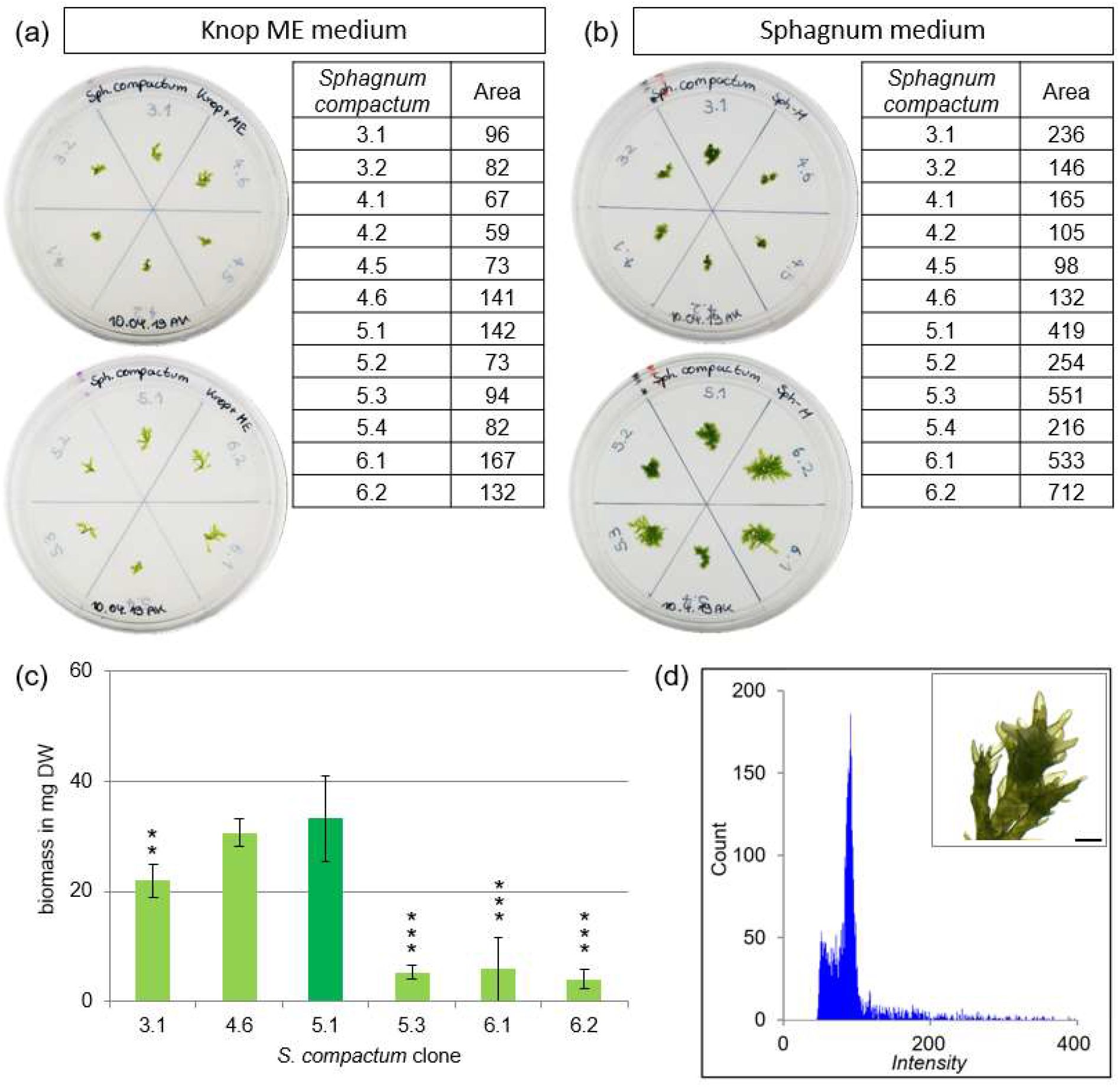
Determination of the best-growing clone of *S. compactum*. Growth determination of 12 *S. compactum* clones on (a) solid Knop ME and (b) solid Sphagnum medium after four weeks of cultivation. The following clones were arranged counterclockwise on the Petri dish: 3.1, 3.2, 4.1, 4.2, 4.5, 4.6 on upper and 5.1, 5.2, 5.3, 5.4, 6.1, 6.2 on lower. Gametophores were cultivated for four weeks. The size of the gametophores was measured on the basis of binary pictures using ImageJ and shown in the table next to it. The six best-growing clones (3.1, 4.6, 5.1, 5.3, 6.1 and 6.2) were selected and the growth was determined in suspension (c) by measuring the dry weight after cultivation of three capitula in flasks containing 50 ml Sphagnum media for six weeks. The y-axis shows the biomass in mg dry weight, the x-axis shows the clone. Data represents mean values with standard deviations of 3 biological replicates (ANOVA p<0.0001). Clone 5.1 yielded significant more biomass compared to the clones 3.1**, 5.3***, 6.1*** and 6.2***. Asterisks represent results of student t-test performed in comparison to clone 5.1. * = p < 0.05, ** = p < 0.01, *** = p < 0.001. The ploidy of the best-growing clone *S. compactum* 5.1 was determined by flow cytometry (FCM). (d) Histograms of gametophore samples measured via FCM after cultivation on solid Sphagnum medium for four months and a picture of *S. compactum* gametophore after four weeks (scale bar = 1 mm). The channel numbers corresponding to the relative fluorescence intensities of the analysed particles is shown on the x-axis, the number of counted events is shown on the y-axis.

**Fig. S6.**
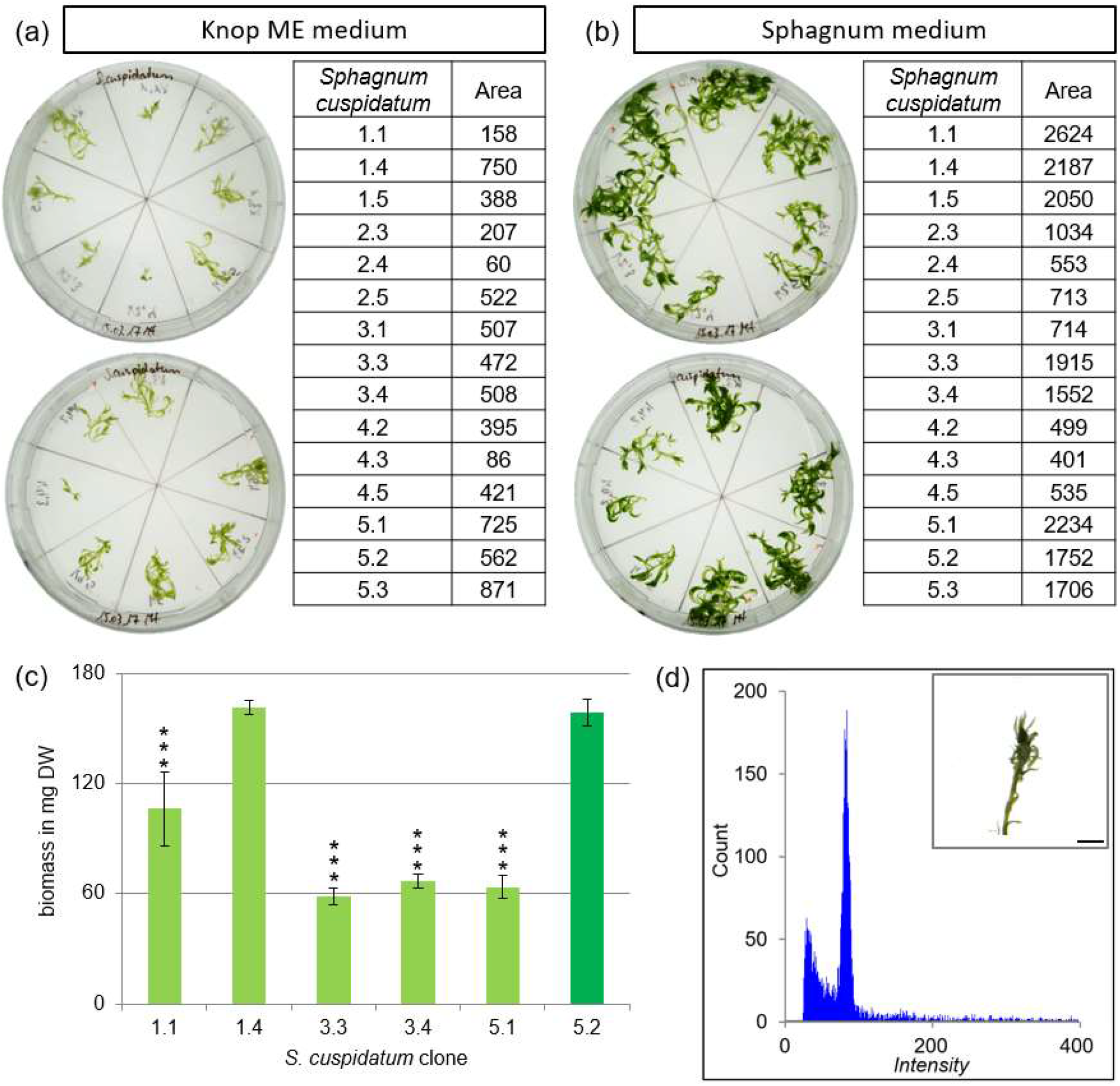
Determination of the best-growing clone of *S. cuspidatum*. Growth determination of 15 *S. cuspidatum* clones on (a) solid Knop ME and (b) solid Sphagnum medium after four weeks of cultivation. The following clones were arranged counterclockwise on the Petri dish: 1.1, 1.4, 1.5, 2.3, 2.4, 2.5, 3.1, 3.3 on upper and 3.4, 4.2, 4.3, 4.5, 5.1, 5.2, 5.3 on lower. Gametophores were cultivated for four weeks. The size of the gametophores was measured on the basis of binary pictures using ImageJ and shown in the table next to it. The six best-growing clones (1.1, 1.4, 3.3, 3.4, 5.1 and 5.2) were selected and the growth was determined in suspension (c) by measuring the dry weight after cultivation of three capitula in flasks containing 50 ml Sphagnum media for six weeks. The y-axis shows the biomass in mg dry weight, the x-axis shows the clone. Data represents mean values with standard deviations of 3 biological replicates, except for clone 1.4 (2 replicates) (ANOVA p<0.0001). Clone 5.2 yielded more biomass compared to the clones 1.1***, 3.3***, 3.4***, 5.1***, but the biomass increase is not significantly better than for clone 5.2. Asterisks represent results of student t-test performed in comparison to clone 2.4. * = p < 0.05, ** = p < 0.01, *** = p < 0.001. The ploidy of the best-growing clone *S. cuspidatum* 5.2 was determined by flow cytometry (FCM). (d) Histograms of gametophore samples measured via FCM after cultivation on solid Sphagnum medium for four months and a picture of *S. cuspidatum* gametophore after four weeks (scale bar = 1 mm). The channel numbers corresponding to the relative fluorescence intensities of the analysed particles is shown on the x-axis, the number of counted events is shown on the y-axis.

**Fig. S7.**
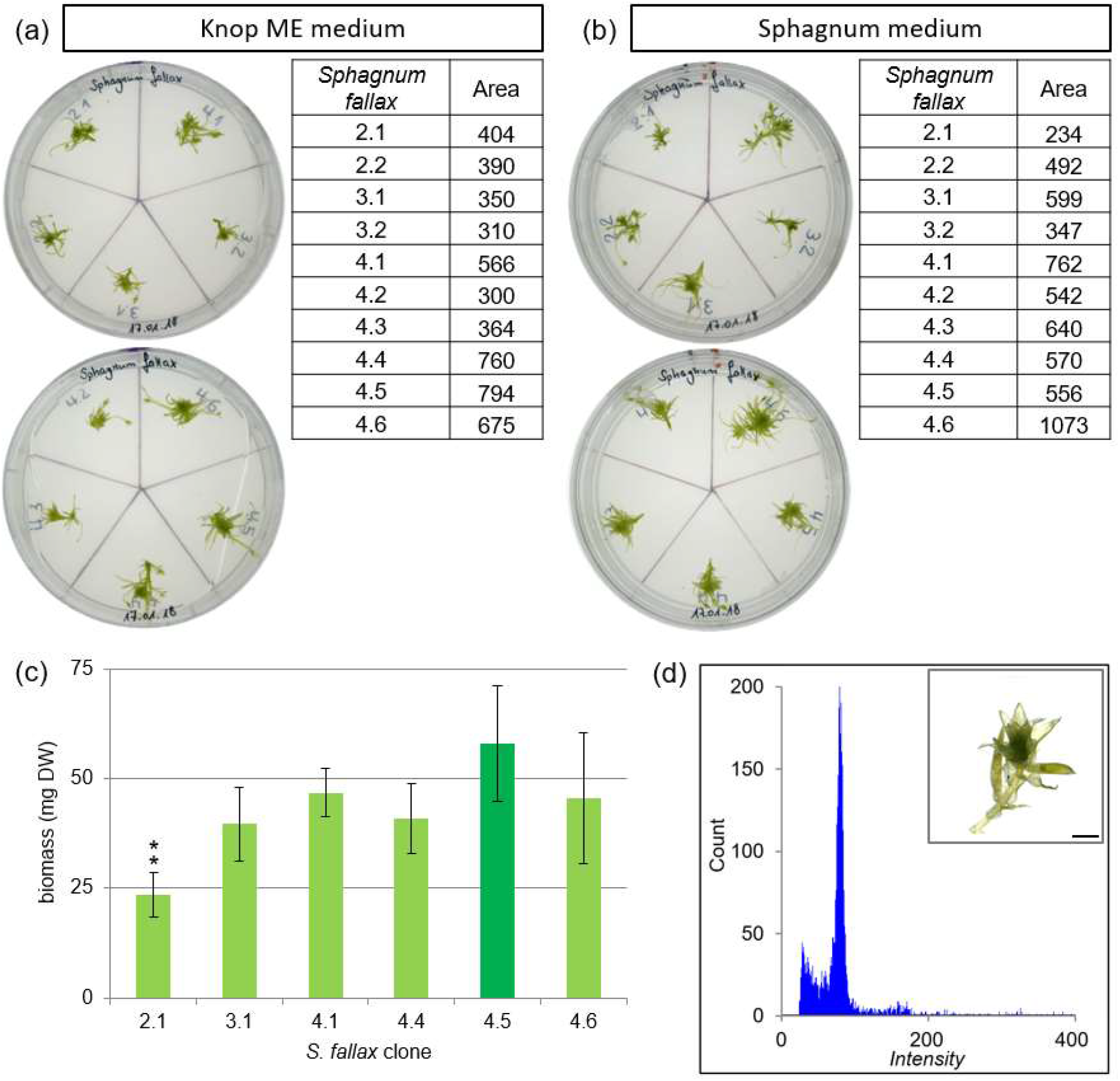
Determination of the best-growing clone of *S. fallax*. Growth determination of 10 *S. fallax* clones on (a) solid Knop ME and (b) solid Sphagnum medium after four weeks of cultivation. The following clones were arranged counter-clockwise on the Petri dish: 2.1, 2.2, 3.1, 3.2, 4.1 on upper and 4.2, 4.3, 4.4, 4.5, 4.6 on lower. Gametophores were cultivated for four weeks. The size of the gametophores was measured on the basis of binary pictures using ImageJ and shown in the table next to it. The six best-growing clones (2.1, 3.1, 4.1, 4.4, 4.5 and 4.6) were selected and the growth was determined in suspension (c) by measuring the dry weight after cultivation of three capitula in flasks containing 50 ml Sphagnum media for six weeks. The y-axis shows the biomass in mg dry weight, the x-axis shows the clone. Data represents mean values with standard deviations of 3 biological replicates, except for clone 4.5 (2 replicates) (ANOVA p<0.05). Clone 4.5 yielded more biomass compared to clone 2.1**, but the biomass increase is not significantly better than for the clones 3.1, 4.1, 4.4 and 4.6. Asterisks represent results of student t-test performed in comparison to clone 4.5. * = p < 0.05, ** = p < 0.01, *** = p < 0.001. The ploidy of the best-growing clone *S. fallax* 4.5 was determined by flow cytometry (FCM). (d) Histograms of gametophore samples measured via FCM after cultivation on solid Sphagnum medium for four months and a picture of *S. fallax* gametophore after four weeks (scale bar = 1 mm). The channel numbers corresponding to the relative fluorescence intensities of the analysed particles is shown on the x-axis, the number of counted events is shown on the y-axis.

**Fig. S8.**
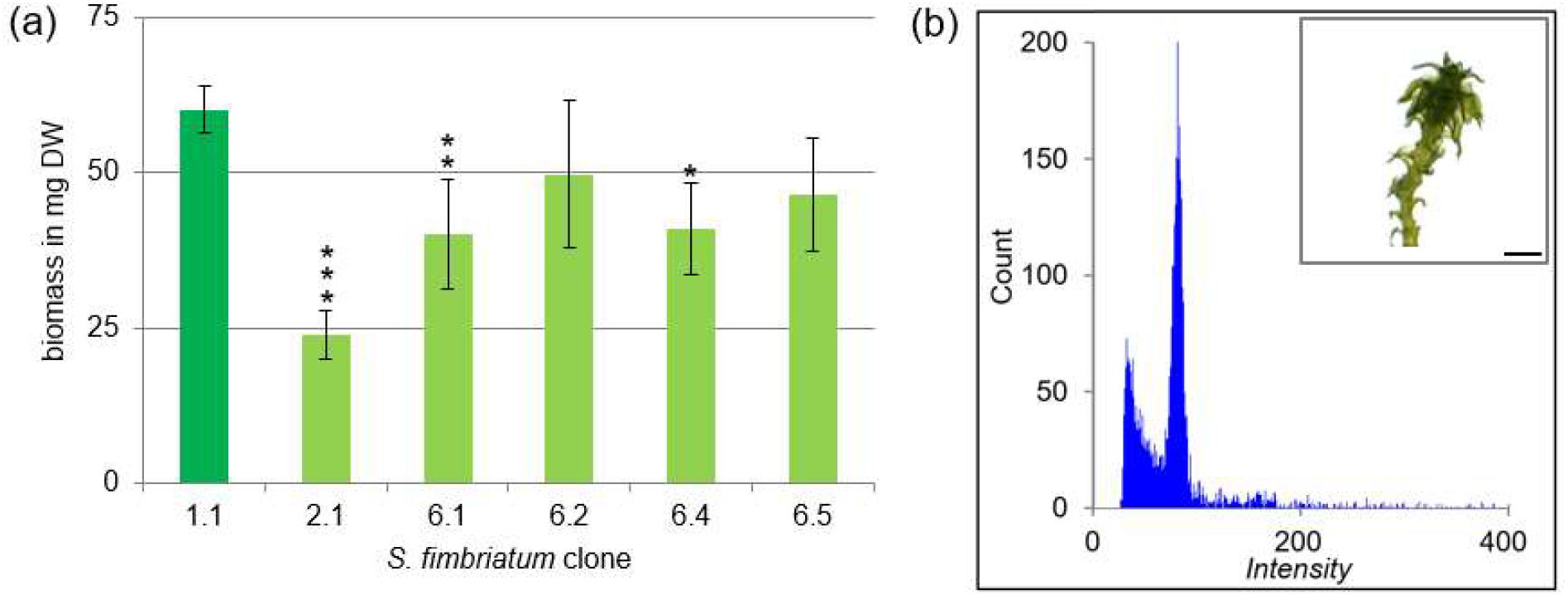
Determination of the best-growing clone of *S. fimbriatum*. The growth of the six clones (1.1, 2.1, 6.1, 6.2, 6.4 and 6.5) was determined in suspension (a) by measuring the dry weight after cultivation of three capitula in flasks containing 50 ml Sphagnum media for six weeks. The y-axis shows the biomass in mg dry weight, the x-axis shows the clone. Data represents mean values with standard deviations of 3 biological replicates (ANOVA p<0.05). Clone 1.1 yielded more biomass compared to the clones 2.1***, 6.1** and 6.4*, but the biomass increase is not significantly better than for clone 6.2 and 6.5. Asterisks represent results of student t-test performed in comparison to clone 1.1. * = p < 0.05, ** = p < 0.01, *** = p < 0.001. The ploidy of the best-growing clone *S. fimbriatum* 1.1 was determined by flow cytometry (FCM). (b) Histograms of gametophore samples measured via FCM after cultivation on solid Sphagnum medium for four months and a picture of *S. fimbriatum* gametophore after four weeks (scale bar = 1 mm). The channel numbers corresponding to the relative fluorescence intensities of the analysed particles is shown on the x-axis, the number of counted events is shown on the y-axis.

**Fig. S9.**
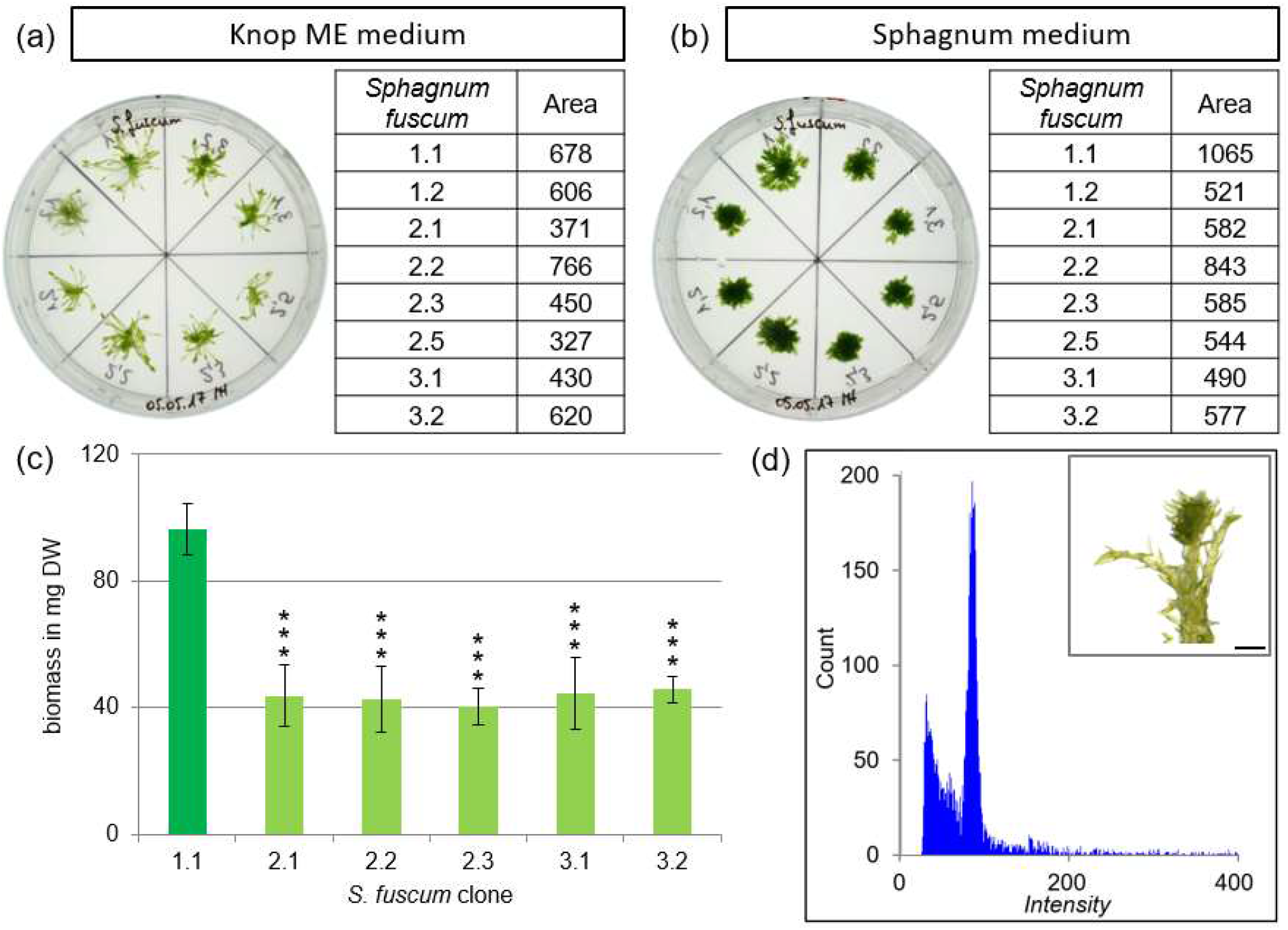
Determination of the best-growing clone of *S. fuscum*. Growth determination of eight *S. fuscum* clones on (a) solid Knop ME and (b) solid Sphagnum medium after four weeks of cultivation. The following clones were arranged counterclockwise on the Petri dish: 1.1, 1.2, 2.1, 2.2, 2.3, 2.5, 3.1, 3.2. Gametophores were cultivated for four weeks. The size of the gametophores was measured on the basis of binary pictures using ImageJ and shown in the table next to it. The six best-growing clones (1.1, 2.1, 2.2, 2.3, 3.1 and 3.2) were selected and the growth was determined in suspension (c) by measuring the dry weight after cultivation of three capitula in flasks containing 50 ml Sphagnum media for six weeks. The y-axis shows the biomass in mg dry weight, the x-axis shows the clone. Data represents mean values with standard deviations of 3 biological replicates (ANOVA p<0.0001). Clone 1.1 yielded significantly more biomass compared to the clones 2.1***, 2.2***, 2.3***, 3.1*** and 3.2***. Asterisks represent results of student t-test performed in comparison to clone 1.1. * = p < 0.05, ** = p < 0.01, *** = p < 0.001. The ploidy of the best-growing clone *S. fuscum* 1.1 was determined by flow cytometry (FCM). (d) Histograms of gametophore samples measured via FCM after cultivation on solid Sphagnum medium for four months and a picture of *S. fuscum* gametophore after four weeks (scale bar = 1 mm). The channel numbers corresponding to the relative fluorescence intensities of the analysed particles is shown on the x-axis, the number of counted events is shown on the y-axis.

**Fig. S10.**
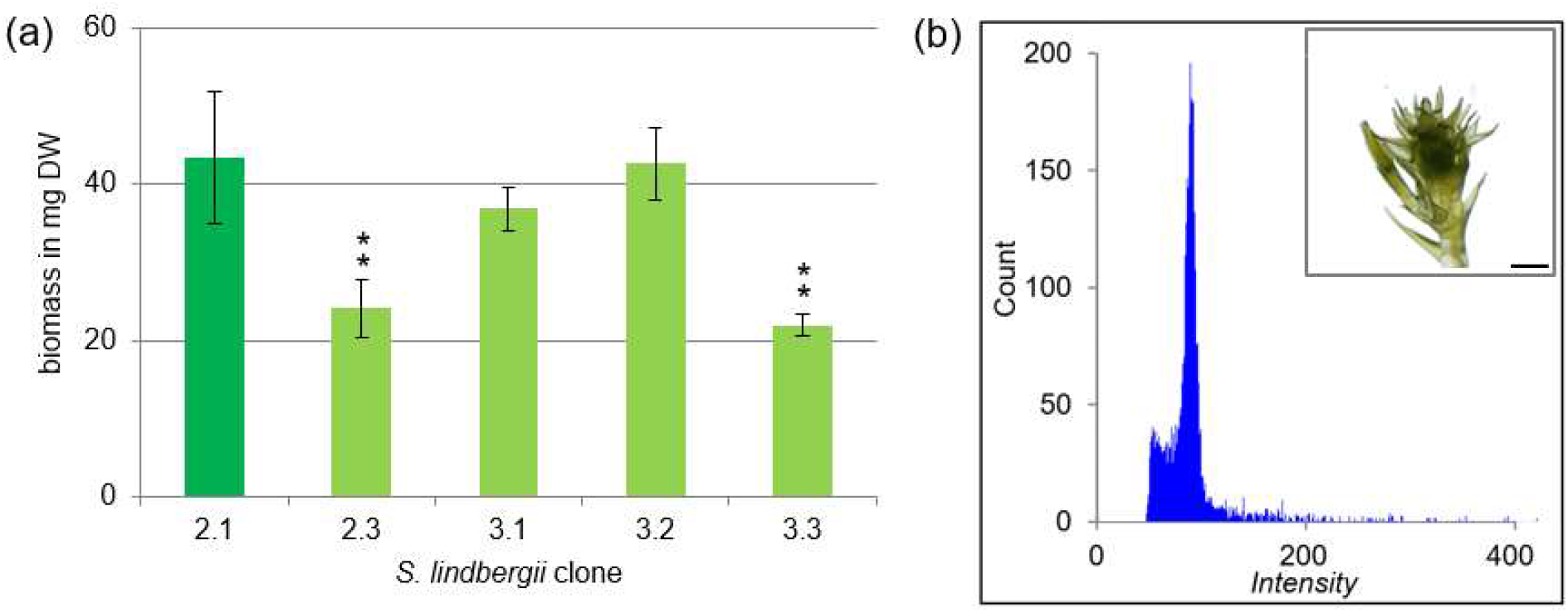
Determination of the best-growing clone of *S. lindbergii*. The growth of the six clones (2.1, 2.3, 3.1, 3.2 and 3.3) was determined in suspension (a) by measuring the dry weight after cultivation of three capitula in flasks containing 50 ml Sphagnum media for six weeks. The y-axis shows the biomass in mg dry weight, the x-axis shows the clone. Data represents mean values with standard deviations of 3 biological replicates, except for clone 3.1 (2 replicates) (ANOVA p<0.05). Clone 2.1 yielded more biomass compared to clone 2.3** and 3.3**, but the biomass increase is not significantly better than for clone 3.1 and 3.2. Asterisks represent results of student t-test performed in comparison to clone 2.1. * = p < 0.05, ** = p < 0.01, *** = p < 0.001. The ploidy of the best-growing clone *S. lindbergii* 2.1 was determined by flow cytometry (FCM). (b) Histograms of gametophore samples measured via FCM after cultivation on solid Sphagnum medium for four months and a picture of *S. lindbergii* gametophore after four weeks (scale bar = 1 mm). The channel numbers corresponding to the relative fluorescence intensities of the analysed particles is shown on the x-axis, the number of counted events is shown on the y-axis.

**Fig. S11.**
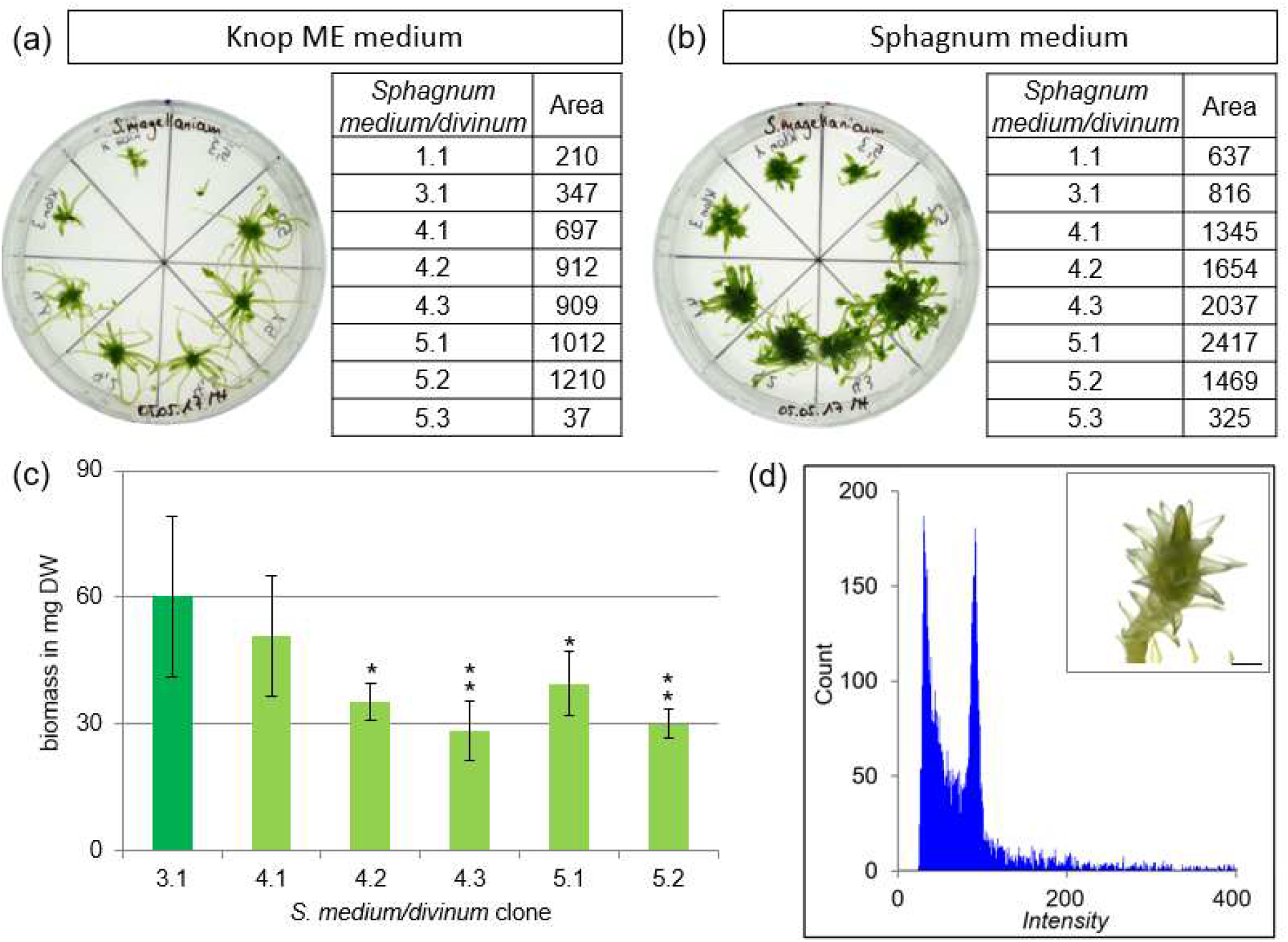
Determination of the best-growing clone of *S. medium/divinum*. Growth determination of eight *S. medium/divinum* clones on (a) solid Knop ME and (b) solid Sphagnum medium after four weeks of cultivation. The following clones were arranged counter-clockwise on the Petri dish: 1.1, 3.1, 4.1, 4.2, 4.3, 5.1, 5.2, 5.3. Gametophores were cultivated for four weeks. The size of the gametophores was measured on the basis of binary pictures using ImageJ and shown in the table next to it. The six best-growing clones (3.1, 4.1, 4.2, 4.3, 5.1 and 5.2) were selected and the growth was determined in suspension (c) by measuring the dry weight after cultivation of three capitula in flasks containing 50 ml Sphagnum media for six weeks. The y-axis shows the biomass in mg dry weight, the x-axis shows the clone. Data represents mean values with standard deviations of 3 biological replicates (ANOVA p<0.05). Clone 3.1 yielded more biomass compared to the clones 4.2*, 4.3**, 5.1* and 5.2**, but the biomass increase is not significantly better than for clone 4.1. Asterisks represent results of student t-test performed in comparison to clone 3.1. * = p < 0.05, ** = p < 0.01, *** = p < 0.001. The ploidy of the best-growing clone *S. medium/divinum* 3.1 was determined by flow cytometry (FCM). (d) Histograms of gametophore samples measured via FCM after cultivation on solid Sphagnum medium for four months and a picture of *S. medium/divinum* gametophore after four weeks (scale bar = 1 mm). The channel numbers corresponding to the relative fluorescence intensities of the analysed particles is shown on the x-axis, the number of counted events is shown on the y-axis.

**Fig. S12.**
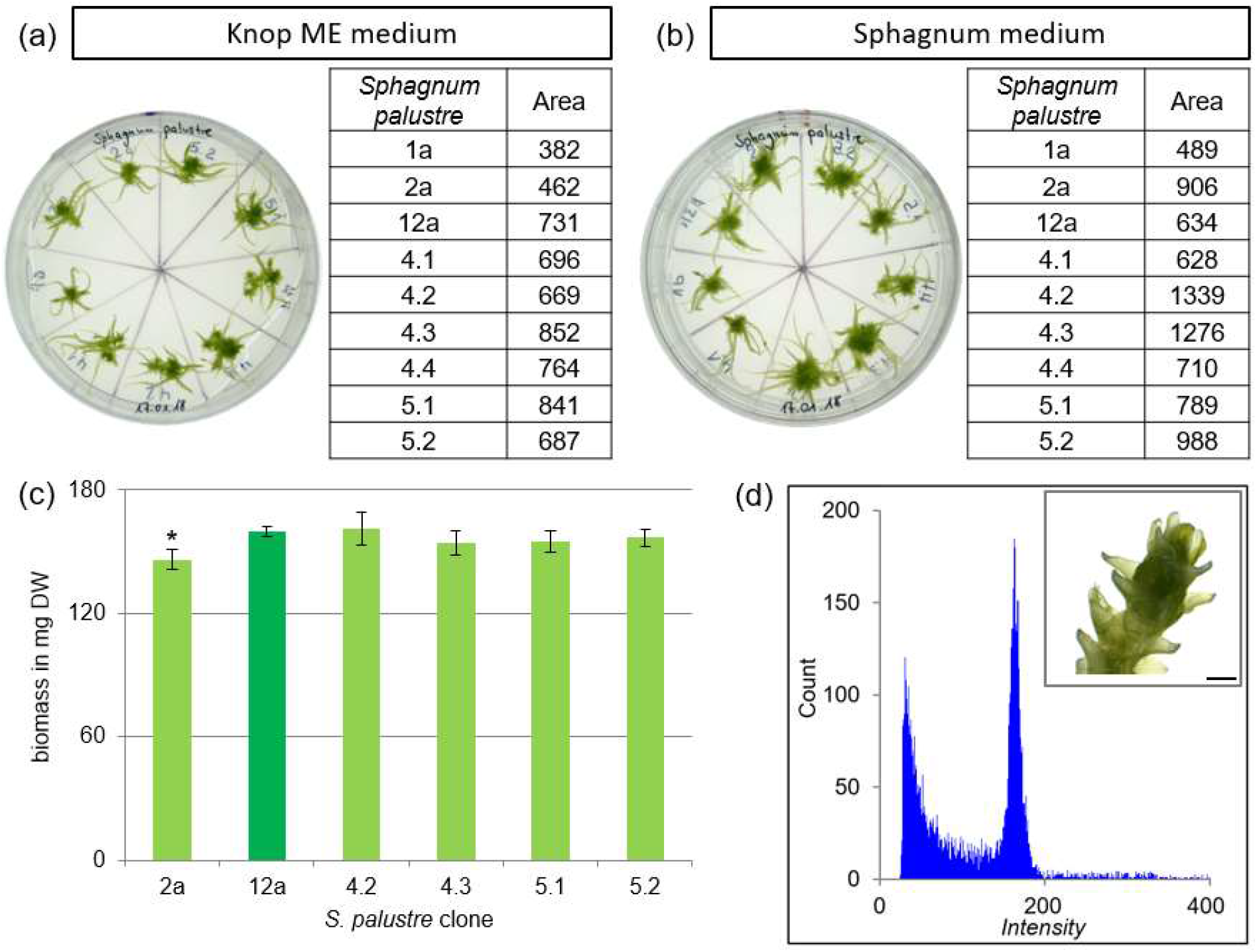
Determination of the best-growing clone of *S. palustre*. Growth determination of nine *S. palustre* clones on (a) solid Knop ME and (b) solid Sphagnum medium after four weeks of cultivation. The following clones were arranged counterclockwise on the Petri dish: 1a, 2a, 12a, 4.1, 4.2, 4.3, 4.4, 5.1, 5.2. Gametophores were cultivated for four weeks. The size of the gametophores was measured on the basis of binary pictures using ImageJ and shown in the table next to it. The six best-growing clones (2a, 12a, 4.2, 4.3, 5.1 and 5.2) were selected and the growth was determined in suspension (c) by measuring the dry weight after cultivation of three capitula in flasks containing 50 ml Sphagnum media for six weeks. The y-axis shows the biomass in mg dry weight, the x-axis shows the clone. Data represents mean values with standard deviations of 3 biological replicates (ANOVA p<0.05). Clone 12a yielded more biomass compared to clone 2a*, but the biomass increase is not significantly better than for the clones 4.2, 4.3, 5.1 and 5.2. Asterisks represent results of student t-test performed in comparison to clone 12a. * = p < 0.05, ** = p < 0.01, *** = p < 0.001. The ploidy of the best-growing clone *S. palustre* 12a was measured by flow cytometry (FCM). (d) Histograms of gametophore samples determined via FCM after cultivation on solid Sphagnum medium for four months and a picture of *S. palustre* gametophore after four weeks (scale bar = 1 mm). The channel numbers corresponding to the relative fluorescence intensities of the analysed particles is shown on the x-axis, the number of counted events is shown on the y-axis.

**Fig. S13.**
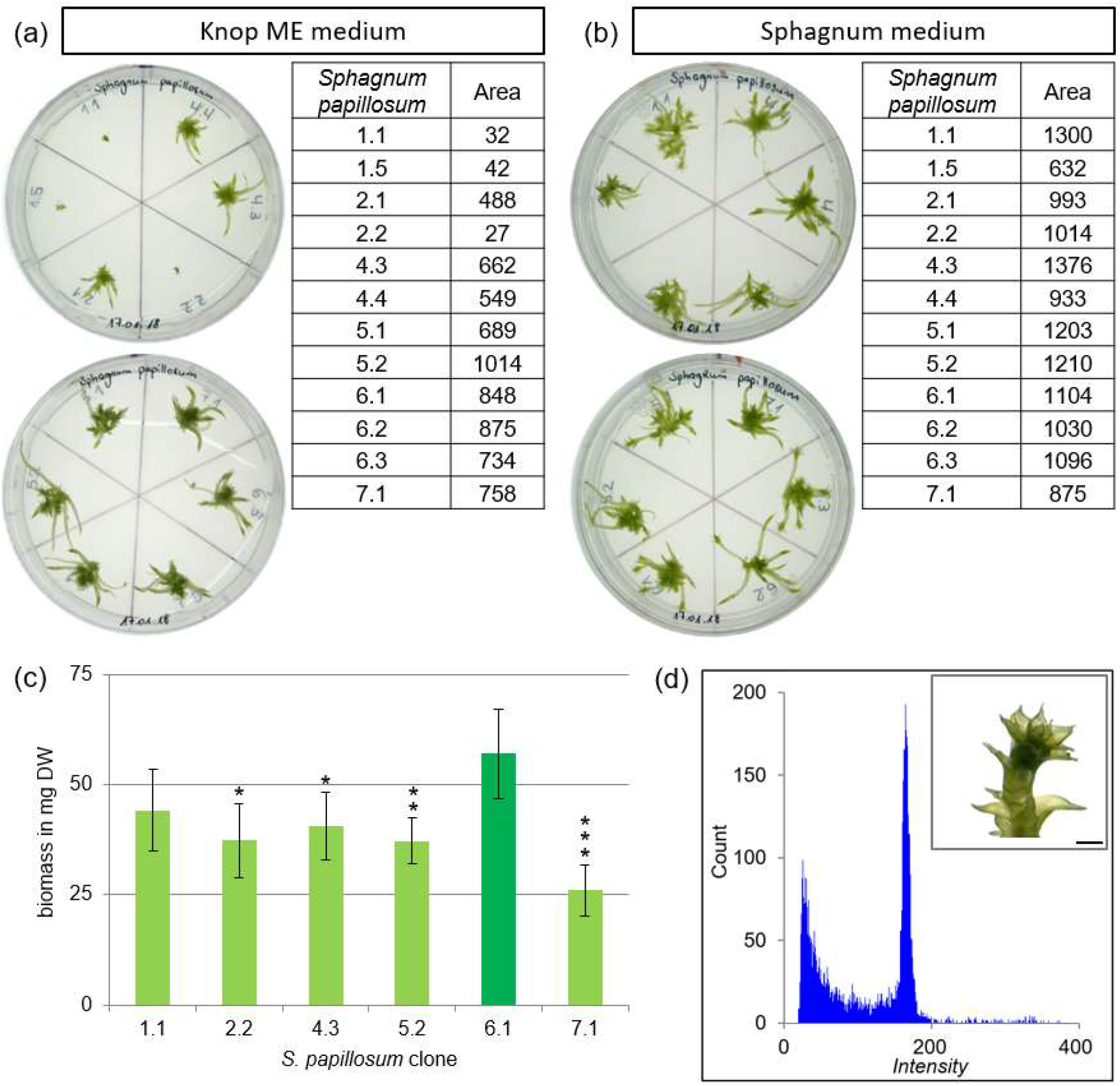
Determination of the best-growing clone of *S. papillosum*. Growth determination of 12 *S. papillosum* clones on (a) solid Knop ME and (b) solid Sphagnum medium after four weeks of cultivation. The following clones were arranged counterclockwise on the Petri dish: 1.1, 1.5, 2.1, 2.2, 4.3, 4.4 on upper and 5.1, 5.2, 6.1, 6.2, 6.3, 7.1 on lower. Gametophores were cultivated for four weeks. The size of the gametophores was measured on the basis of binary pictures using ImageJ and shown in the table next to it. The six best-growing clones (1.1, 2.2, 4.3, 5.2, 6.1 and 7.1) were selected and the growth was determined in suspension (c) by measuring the dry weight after cultivation of three capitula in flasks containing 50 ml Sphagnum media for six weeks. The y-axis shows the biomass in mg dry weight, the x-axis shows the clone. Data represents mean values with standard deviations of 3 biological replicates (ANOVA p<0.05). Clone 6.1 yielded more biomass compared to the clones 2.2*, 4.3*, 5.2**, and 7.1***, but the biomass increase is not significantly better than for clone 1.1. Asterisks represent results of student t-test performed in comparison to clone 6.1. * = p < 0.05, ** = p < 0.01, *** = p < 0.001. The ploidy of the best-growing clone *S. papillosum* 6.1 was determined by flow cytometry (FCM). (d) Histograms of gametophore samples measured via FCM after cultivation on solid Sphagnum medium for four months and a picture of *S. papillosum* gametophore after four weeks (scale bar = 1 mm). The channel numbers corresponding to the relative fluorescence intensities of the analysed particles is shown on the x-axis, the number of counted events is shown on the y-axis.

**Fig. S14.**
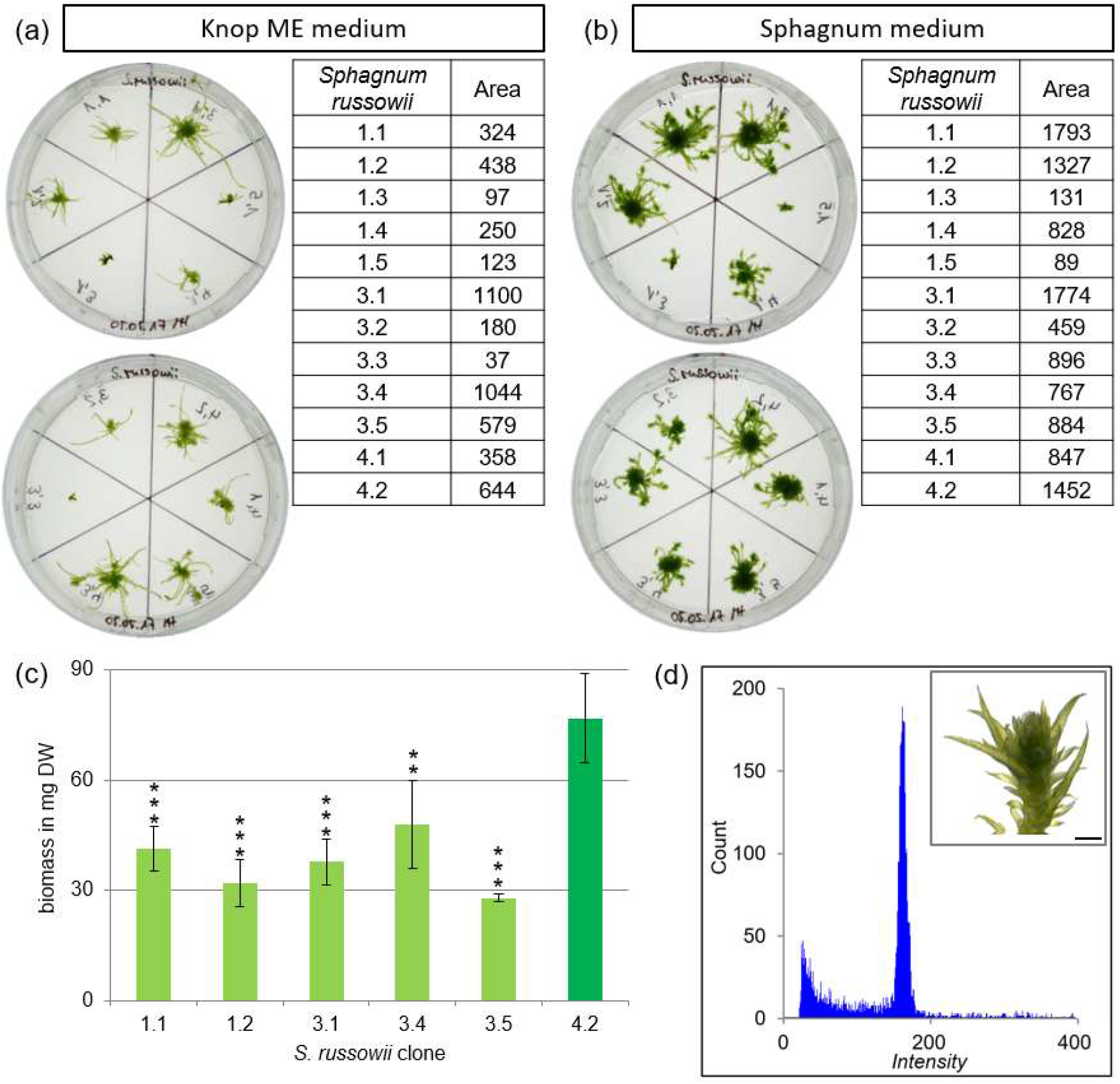
Determination of the best-growing clone of *S. russowii*. Growth determination of 12 *S. russowii* clones on (a) solid Knop ME and (b) solid Sphagnum medium after four weeks of cultivation. The following clones were arranged counterclockwise on the Petri dish: 1.1, 1.2, 1.3, 1.4, 1.5, 3.1 on upper and 3.2, 3.3, 3.4, 3.5, 4.1, 4.2 on lower. Gametophores were cultivated for four weeks. The size of the gametophores was measured on the basis of binary pictures using ImageJ and shown in the table next to it. The six best-growing clones (1.1, 1.2, 3.1, 3.4, 3.5 and 4.2) were selected and the growth was determined in suspension (c) by measuring the dry weight after cultivation of three capitula in flasks containing 50 ml Sphagnum media for six weeks. The y-axis shows the biomass in mg dry weight, the x-axis shows the clone. Data represents mean values with standard deviations of 3 biological replicates (ANOVA p=0.0001). Clone 4.2 yielded significantly more biomass compared to the clones 1.1***, 1.2***, 3.1***, 3.4** and 3.5***. Asterisks represent results of student t-test performed in comparison to clone 4.2. * = p < 0.05, ** = p < 0.01, *** = p < 0.001. The ploidy of the best-growing clone *S. russowii* 4.2 was determined by flow cytometry (FCM). (d) Histograms of gametophore samples measured via FCM after cultivation on solid Sphagnum medium for four months and a picture of *S. russowii* gametophore after four weeks (scale bar = 1 mm). The channel numbers corresponding to the relative fluorescence intensities of the analysed particles is shown on the x-axis, the number of counted events is shown on the y-axis.

**Fig. S15.**
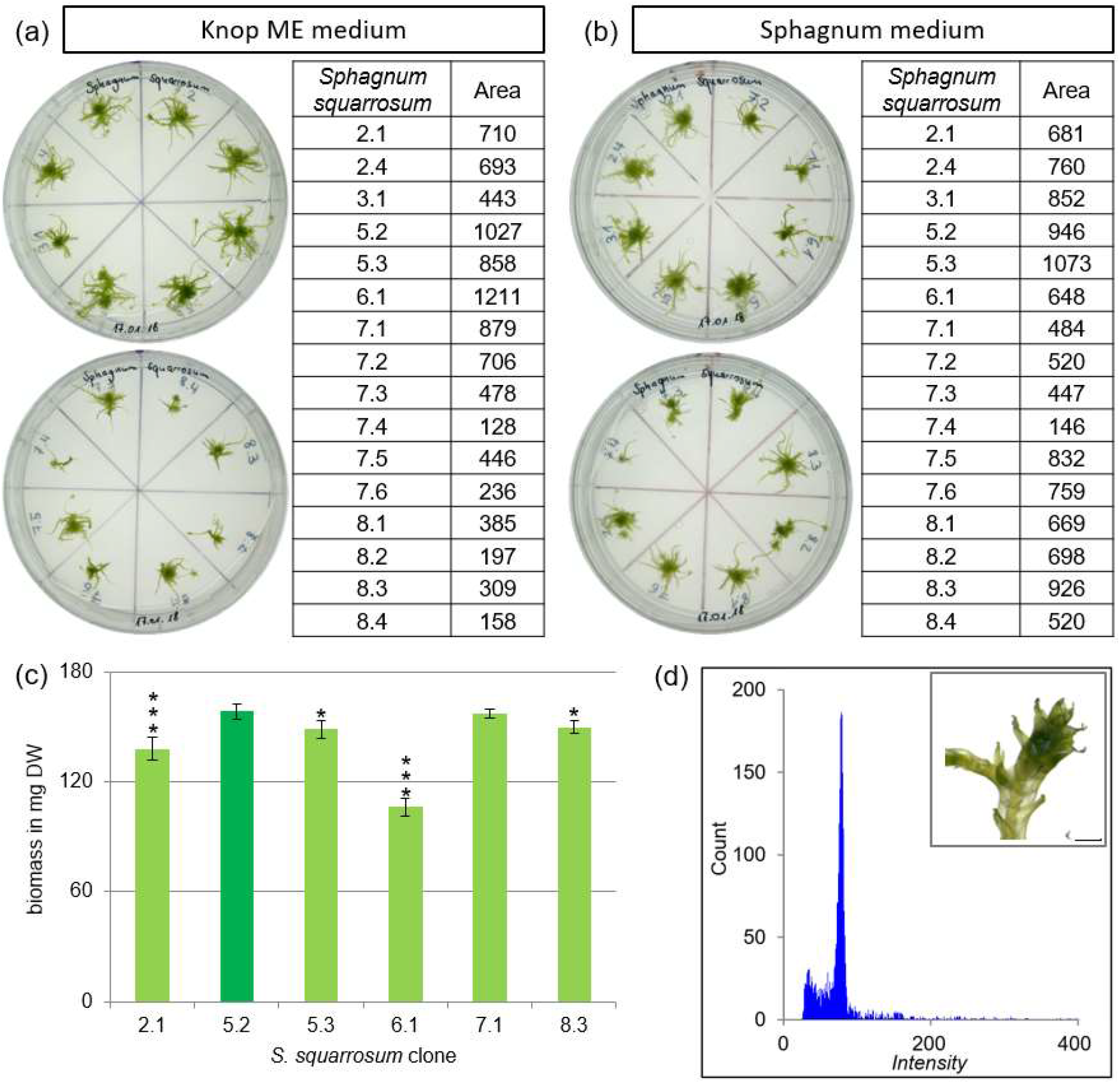
Determination of the best-growing clone of *S. squarrosum*. Growth determination of 16 *S. squarrosum* clones on (a) solid Knop ME and (b) solid Sphagnum medium after four weeks of cultivation. The following clones were arranged counterclockwise on the Petri dish: 2.1, 2.4, 3.1, 5.2, 5.3, 6.1, 7.1, 7.2 on upper and 7.3, 7.4, 7.5, 7.6, 8.1, 8.2, 8.3, 8.4 on lower. Gametophores were cultivated for four weeks. The size of the gametophores was measured on the basis of binary pictures using ImageJ and shown in the table next to it. The six best-growing clones (2.1, 5.2, 5.3, 6.1, 7.1 and 8.3) were selected and the growth was determined in suspension (c) by measuring the dry weight after cultivation of three capitula in flasks containing 50 ml Sphagnum media for six weeks. The y-axis shows the biomass in mg dry weight, the x-axis shows the clone. Data represents mean values with standard deviations of 3 biological replicates (ANOVA p<0.0001). Clone 5.2 yielded more biomass compared to the clones 2.1***, 5.3*, 6.1*** and 8.3*, but the biomass increase is not significantly better than for clone 7.1. Asterisks represent results of student t-test performed in comparison to clone 5.2. * = p < 0.05, ** = p < 0.01, *** = p < 0.001. The ploidy of the best-growing clone *S. squarrosum* 5.2 was determined by flow cytometry (FCM). (d) Histograms of gametophore samples measured via FCM after cultivation on solid Sphagnum medium for four months and a picture of *S. squarrosum* gametophore after four weeks (scale bar = 1 mm). The channel numbers corresponding to the relative fluorescence intensities of the analysed particles is shown on the x-axis, the number of counted events is shown on the y-axis.

**Fig. S16.**
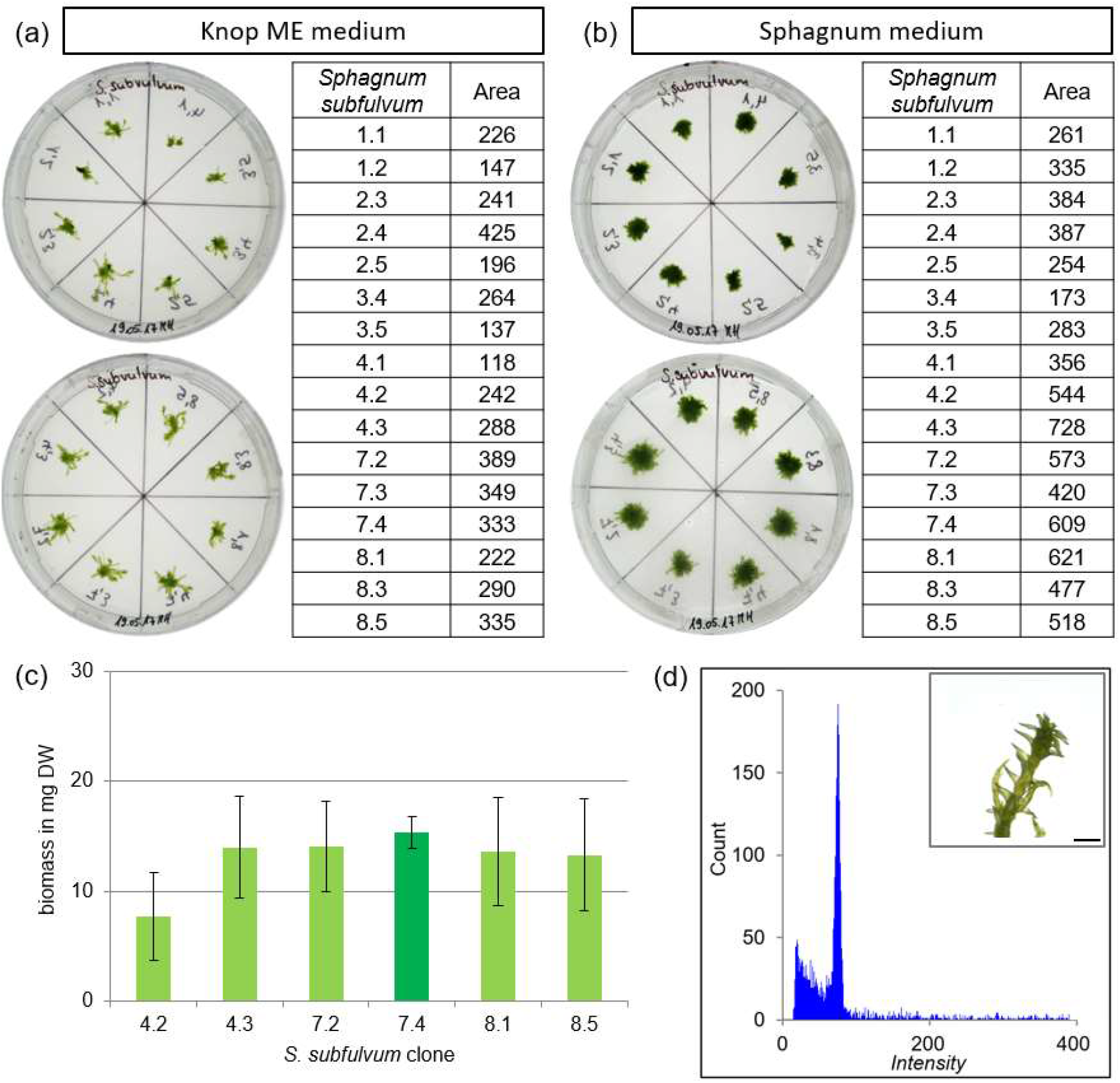
Determination of the best-growing clone of *S. subfulvum*. Growth determination of 16 *S. subfulvum* clones on (a) solid Knop ME and (b) solid Sphagnum medium after four weeks of cultivation. The following clones were arranged counterclockwise on the Petri dish: 1,1. 1.2, 2.3, 2.4, 2.5, 3.4, 3.5, 4.1 on upper and 4.2, 4.3, 7.2, 7.3, 7.4, 8.1, 8.3, 8.5 on lower. Gametophores were cultivated for four weeks. The size of the gametophores was measured on the basis of binary pictures using ImageJ and shown in the table next to it. The six best-growing clones (4.2, 4.3, 7.2, 7.4, 8.1 and 8.5) were selected and the growth was determined in suspension (c) by measuring the dry weight after cultivation of three capitula in flasks containing 50 ml Sphagnum media for six weeks. The y-axis shows the biomass in mg dry weight, the x-axis shows the clone. Data represents mean values with standard deviations of 3 biological replicates (ANOVA p>0.05). Clone 7.4 was selected as best-growing clone. The ploidy of the best-growing clone *S. subfulvum* 7.4 was determined by flow cytometry (FCM). (d) Histograms of gametophore samples measured via FCM after cultivation on solid Sphagnum medium for four months and a picture of *S. subfulvum* gametophore after four weeks (scale bar = 1 mm). The channel numbers corresponding to the relative fluorescence intensities of the analysed particles is shown on the x-axis, the number of counted events is shown on the y-axis.

**Fig. S17.**
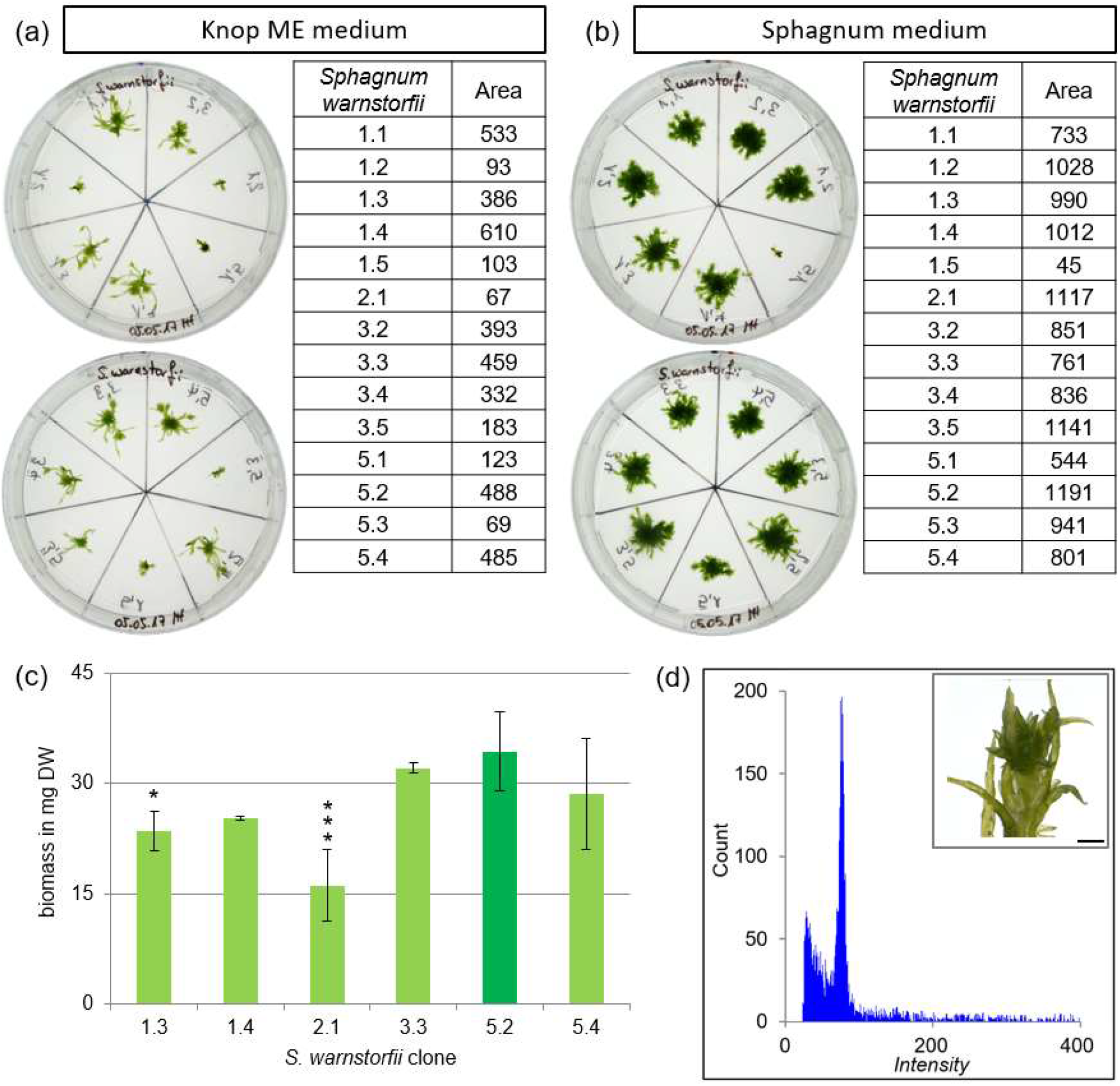
Determination of the best-growing clone of *S. warnstorfii*. Growth determination of 14 *S. warnstorfii* clones on (a) solid Knop ME and (b) solid Sphagnum medium after four weeks of cultivation. The following clones were arranged counterclockwise on the Petri dish: 1,1. 1.2, 1.3, 1.4, 1.5, 2.1, 3.2 on upper and 3.3, 3.4, 3.5, 5.1, 5.2, 5.3, 5.4 on lower. Gametophores were cultivated for four weeks. The size of the gametophores was measured on the basis of binary pictures using ImageJ and shown in the table next to it. The six best-growing clones (4.2, 4.3, 7.2, 7.4, 8.1 and 8.5) were selected and the growth was determined in suspension (c) by measuring the dry weight after cultivation of three capitula in flasks containing 50 ml Sphagnum media for six weeks. The y-axis shows the biomass in mg dry weight, the x-axis shows the clone. Data represents mean values with standard deviations of 3 biological replicates, except for clone 1.4 and 3.3 (2 replicates) (ANOVA p<0.05). Clone 5.2 yielded more biomass compared to clone 1.3* and 2.1***, but the biomass increase is not significantly better than for the clones 1.4, 3.3 and 5.4. Asterisks represent results of student t-test performed in comparison to clone 5.2. * = p < 0.05, ** = p < 0.01, *** = p < 0.001. The ploidy of the best-growing clone *S. warnstorfii* 5.2 was determined by flow cytometry (FCM). (d) Histograms of gametophore samples measured via FCM after cultivation on solid Sphagnum medium for four months and a picture of *S. warnstorfii* gametophore after four weeks (scale bar = 1 mm). The channel numbers corresponding to the relative fluorescence intensities of the analysed particles is shown on the x-axis, the number of counted events is shown on the y-axis.

